# A compendium of Amplification-Related Gain Of Sensitivity (ARGOS) genes in human cancer

**DOI:** 10.1101/2023.12.16.571980

**Authors:** Veronica Rendo, Michael Schubert, Nicholas Khuu, Maria F Suarez Peredo Rodriguez, Kaimeng Huang, Michelle Swift, Yizhou He, Johanna Zerbib, Ross Smith, Jonne Raaijmakers, Pratiti Bandopadhayay, Lillian M. Guenther, Justin H. Hwang, Amanda Iniguez, Susan Moody, Ji-Heui Seo, Elizabeth Stover, Levi Garraway, William C. Hahn, Kimberly Stegmaier, René H. Medema, Dipanjan Chowdhury, Maria Colomé-Tatché, Uri Ben-David, Rameen Beroukhim, Floris Foijer

**Affiliations:** Department of Medical Oncology and Center for Neuro-Oncology, Dana-Farber Cancer Institute, Boston, MA, USA; Department of Cancer Biology, Dana-Farber Cancer Institute, Boston, MA, USA; Harvard Medical School, Boston, MA, USA; Broad Institute of Harvard and MIT, Cambridge, MA, USA; Department of Immunology, Genetics and Pathology, Uppsala University, Uppsala, Sweden; Oncode Institute, Division of Cell Biology, The Netherlands Cancer Institute, Amsterdam, Netherlands; European Research Institute for the Biology of Ageing, University Medical Center Groningen, Groningen, Netherlands; Institute of Computational Biology, Helmholtz Munich, Neuherberg, Germany; Department of Radiation Oncology, Dana-Farber Cancer Institute, Boston, MA, USA; Department of Human Molecular Genetics & Biochemistry, Tel Aviv University, Tel Aviv, Israel; Department of Pediatrics, Dana-Farber Cancer Institute, Boston, MA, USA; St. Jude Children’s Research Hospital, Department of Oncology, Memphis, TN, USA; Division of Hematology, Oncology, and Transplantation, University of Minnesota, Minneapolis, MN, USA; Moonlight Bio, Seattle, WA, USA; Department of Cancer Biology, Perelman School of Medicine at the University of Pennsylvania, Philadelphia, PA, USA; Biomedical Center (BMC), Physiological Chemistry, Ludwig Maximilians University, Munich, Germany

## Abstract

Chromosomal gains are among the most frequent somatic genetic alterations occurring in cancer. While the effect of sustained oncogene expression has been characterized, the impact of copy-number gains affecting collaterally-amplified “bystander” genes on cellular fitness remains less understood. To investigate this, we built a comprehensive map of dosage compensations across human cancers by integrating expression and copy number profiles from over 8,000 TCGA tumors and CCLE cell lines. Further, we analyzed the effect of gene overexpression across 17 human cancer ORF screens to provide an overview of genes that prove toxic to cancer cells when overexpressed. Combining these two independent approaches we propose a class of ‘Amplification-Related Gain Of Sensitivity’ (ARGOS) genes. These genes are located in commonly amplified regions of the genome, have lower expression levels than expected by their copy-number status, and are toxic to cancer cells when overexpressed. We experimentally validated *CDKN1A* and *RBM14* as high-confidence pan-cancer ARGOS genes in lung and breast cancer cell line models. We additionally suggest that RBM14’s mechanism of toxicity involves altered DNA damage response and innate immune signaling processes following gene overexpression. Finally, we provide a comprehensive catalog of compensated, toxic, and ARGOS genes as a community resource.

## Introduction

Due to genomic instability, human cancers accumulate somatic mutations over time. The most frequent type of genomic alterations are copy number changes, affecting on average ∼30% of a tumor’s genome ^1,2^. Somatic copy number alterations (sCNAs) can target focal regions of the genome (e.g. amplification of the oncogene *MYC* in chromosome 8q, or deletion of the tumor suppressor *RB1* in chromosome 13q), but often comprise chromosome arm-level events that span hundreds of collaterally-altered genes located in proximity to the cancer driver genes. Such large-scale events shape the transcriptional and translational profile of tumor cells, as chromosomal copy number changes may affect genes with essential roles in tumor progression and viability ^3^.

In human cancers, changes in DNA copy number tend to be tightly correlated with mRNA expression levels ^4,5^. However, uncoupling of gene expression from DNA copy number has been described by multiple mechanisms at the genomic (e.g. rearrangements), epigenetic (e.g. promoter hypermethylation) and post-translational (e.g. buffering copy number imbalances in protein complex members) levels ^6–12^. These discrepancies between gene expression levels and copy number suggest that such large-scale changes may have a negative impact on cellular fitness. Such alterations may also create new dependencies. For example, loss of ‘Copy number alterations Yielding Cancer Liabilities Owing to Partial losS’ (CYCLOPS) genes renders cells dependent on the remaining copy ^13^. Similarly, loss of heterozygosity events spanning passenger metabolic and essential genes, as well as homozygous losses of gene paralogs, create unique dependencies in tumor cells that can be exploited therapeutically ^14–19^.

In this study, we sought to investigate whether copy number gains can also become cancer liabilities, as they affect the expression of multiple genes with diverse biological functions and impact cellular fitness. Oncogene overexpression is known to cause oncogene-induced senescence, a tumor suppressive state characterized by an arrest in cell growth ^20–22^. However, the impact of copy number gains affecting “bystander” genes, co-amplified with oncogenes, is less understood. We hypothesized that some genes located in commonly amplified regions of the genome could be toxic to the cell when overexpressed upon gain, triggering mechanisms of gene compensation to attenuate their overexpression. For this new class of ‘Amplification-Related Gain Of Sensitivity’ (ARGOS) genes, we sought to investigate how overexpressed genes may become tumor-toxic and the compensation mechanisms that cells acquire to abrogate this effect.

## Results

### Copy-number changes often affect non-driver genes whose expression is nevertheless altered

Much of the genome is frequently gained or lost in cancer (**Fig. 1a**). This is because individual copy-number alterations, selected by cancer driver events, typically affect large sections of the genome (**Fig. 1b**). Although Oncogenes (OGs) are more frequently amplified and less frequently lost, and Tumor Suppressor genes (TSGs) show the opposite trend (**Fig. S1a-b**), most of the frequently amplified genes are neither OGs nor TSGs, but likely “collaterally altered” by the gain/loss of large chromosomal regions (**Fig. S1c-d**).

**Fig. 1.**
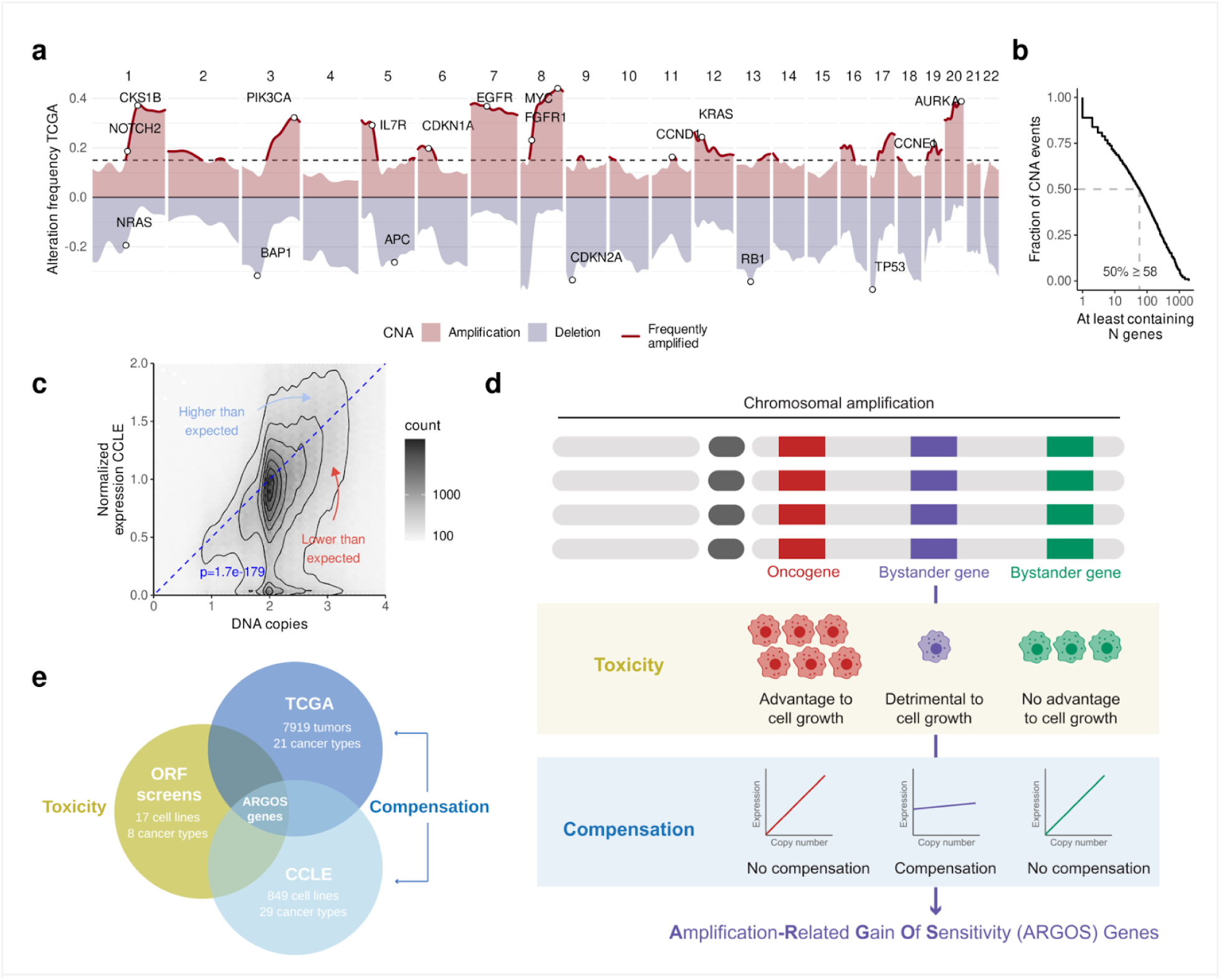
Approach to identify ARGOS genes. **(a)** The landscape of CNAs with the frequency of amplifications and deletions across TCGA tumors shows preferential gains of OGs and losses of TSGs. Genes gained in over 15% of samples (dotted line) were considered commonly amplified. **(b)** Individual CNA events typically contain multiple genes at a median of 58. **(c)** RNA expression scales with DNA copy number in the CCLE, although there is a considerable spread around this trend. We refer to genes expressed consistently lower or higher than the expectation as compensated and hyperactivated, respectively. **(d)** ARGOS genes are identified by genes that are collaterally affected by amplifications, which are also detrimental to cell growth and show compensation upon copy number gains. **(e)** In practical terms, we employ the CCLE and TCGA cohorts to identify compensated genes and confirm the toxicity phenotype by repurposing previously published ORF screens.

**Fig. S1.**
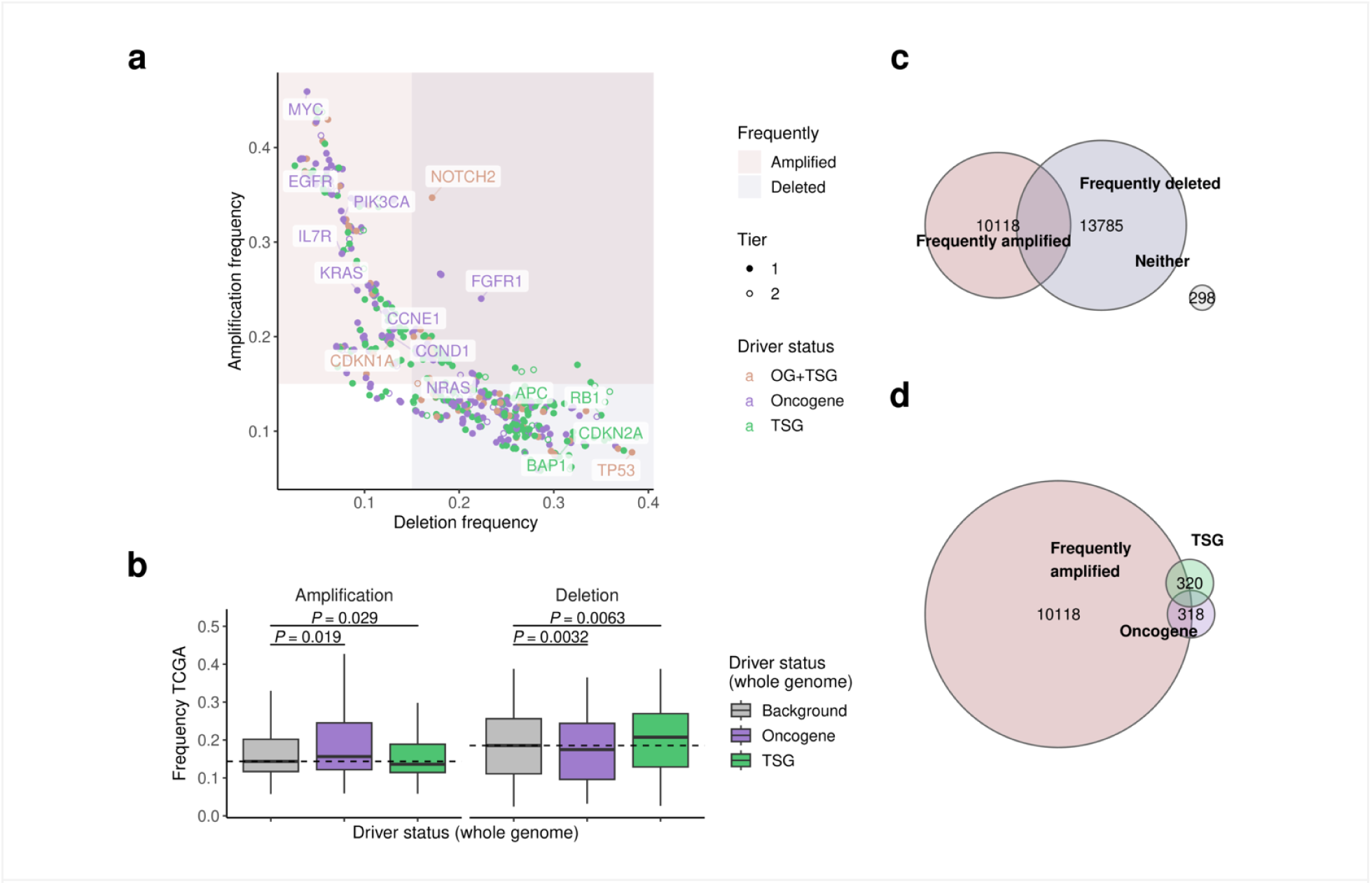
Landscape of copy number alterations. **(a)** Frequency of amplifications and deletions in the TCGA with COSMIC driver status highlighted. **(b)** OGs are more frequently amplified, less frequently deleted and TSGs the opposite, **(c)** however most of the cancer genome is frequently amplified or deleted, and **(d)** most amplifications and deletions are not OGs or TSGs.

Nevertheless, the expression levels of the vast majority of genes, including these “bystander” genes, scale with their copy number ^4,23^. This is true across human tumors ^4^, across cell lines in the Cancer Cell Line Encyclopedia (CCLE) (**Fig. 1c**), and also in isogenic RPE-1 cells with individual gained chromosomes ^24^. However, we also observe considerable variation around this trend (**Fig. 1c**). If amplified genes are consistently expressed lower or higher than expected we refer to them as “compensated” or “hyperactivated”, respectively. We were especially interested in compensated genes because these might reflect negative selective pressures resulting from fitness decreases due to amplification-driven overexpression. That is, genes whose overexpression is “toxic” to cancer cells might exhibit substantial compensation when amplified.

To more directly assess gene toxicity, we assembled gene sensitivity data from 17 different Open Reading Frame (ORF) overexpression screens performed across 10 tumor types ^25–33^. We then quantified which genes strongly decreased in abundance across these screens without additional selection pressure. Combining these toxic genes with genes that we identified as “compensated”, we aimed to identify the set of ARGOS genes whose amplification could jeopardize cancer cell fitness (**Fig. 1d-e**).

### Genes are consistently compensated in CCLE and TCGA

We developed a computational method to detect genes that are consistently compensated in their expression relative to their copy number, both across (pan-cancer) and for individual cancer types (tissue-specific). First, we selected samples with copy-neutral and amplified genes from both large human cancer cell line (CCLE ^34^) and tumor (TCGA ^35^) cohorts. We then built a Bayesian Negative Binomial regression model between these two variables, including a variable for copy number vs. gene expression scaling and one for its deviation (**Fig. 2a**). However, in human tumors, non-cancer cells can represent a large fraction of the cells in a sample (reflecting low tumor purity). These impurities would be expected to modify observed expression levels relative to the expression levels within the cancer cells. To account for this, we explicitly modeled cancer and non-cancer contributions to the observed gene expression in TCGA. This provided us with a compensation score where − 1 indicates complete compensation for cancer cells (*i.e.*, no expression changes with amplifications) and + 1 indicates full hyperactivation (*i.e.*, twice the gene expression that we would expect based on its purity-corrected copy number change) (**Supp. Table 1-2**).

**Fig. 2.**
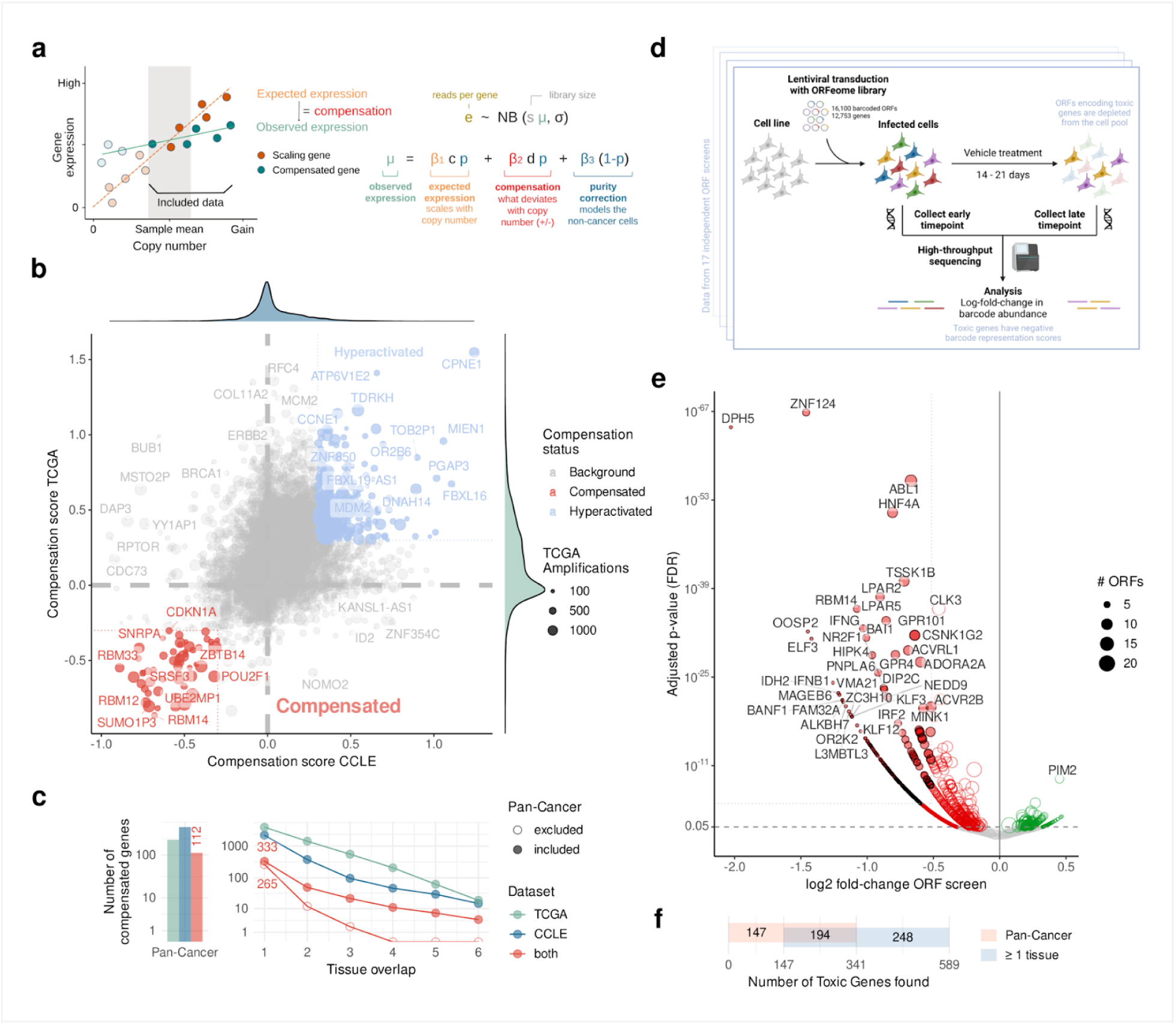
Compensated and toxic genes. **(a)** Using a Bayesian Negative Binomial regression model, we split the expression of each gene in CCLE and TCGA into components that are scaling (orange) and deviating (red) from DNA content. For TCGA data, we explicitly take into account non-cancer cells (blue). We apply this model across (pan-cancer) and for individual cancer types (tissue-specific). **(b)** In the pan-cancer model, deregulation is well correlated between CCLE and TCGA, and we identify commonly compensated (red) and hyperactivated genes (blue). **(c)** Number of pan-cancer and tissue-specific compensated genes, specific to either CCLE or TCGA datasets or common to both. Tissue overlap indicates genes identified in multiple tissue-specific models. Numbers mentioned in-text are highlighted. **(d)** We utilize ORF overexpression screens to quantify barcode abundance for outgrowth after the selection marker, which **(e)** identifies genes that are promoting (green) or attenuating (red) cell growth when overexpressed. Genes passing the significance threshold are shown with a black outline. **(f)** Numbers of pan-cancer vs. tissue-specific toxic (attenuating) genes and their overlap.

As expected, most genes scaled with copy number in both the CCLE and TCGA (pan-cancer analysis; **Fig. 2b**). Compensation scores were significantly correlated between the two datasets, especially when explicitly controlling for non-cancer cells in TCGA data (*P* < 10^-300^, R^2^ = 0.09; **Fig. S2a**). We did not observe an overall difference in compensation scores between genes that are commonly amplified, deleted, or copy-neutral (**Fig. S2c**); or for OGs, TSGs, and genes that are neither (**Fig. S2d**). However, we found splicing and RNA processing genes (Gene Ontology) consistently compensated over the expected scaling, whereas DNA replication and repair genes were hyperactivated when amplified (**Fig. S2b**).

Numbers of pan-cancer vs. tissue-specific toxic (attenuating) genes and their overlap.Using a cutoff of 30% less expression than expected, we identified 112 genes to be compensated in both CCLE and TCGA across cancer types (**Fig. 2b-c**). Among these, we found nine members of the hnRNP family (heterogeneous ribonucleoprotein particle), seven RPLs (Ribosomal protein L), five RBMs (RNA-binding motif), four SRSFs (arginine/serine-rich splicing factor), and three ZNF (Zinc finger) genes. Five are listed as OGs in the COSMIC database ^36^ (*CHD4*, *DGCR8*, *EWSR1*, *HNRNPA2B1*, *SRSF3*), four as TSGs (*CHD2*, *CTCF*, *FUS*, *SFQP*), and two are labeled as both (*CDKN1A* and *DDB2*).

To validate these as compensated genes, we examined gene expression and calculated gene compensation scores in isogenic RPE-1 clones with gained chromosomes (either chromosome 7 or a combination of chromosome 7 and 22; or 8, 9 and 18; for 7, 10 and 7 compensated genes **Fig. S2e**) ^10,37^. We confirmed that the compensated genes residing on the respective gained chromosomes were expressed less than non-compensated genes thereon (*P* < 0.05). Furthermore, our compensated genes showed significantly higher associations between genomic gain and diseases other than cancer compared to non-compensated genes (“Triplosensitivity”; **Fig. S2f**) ^38^. Combining the evidence of these four independent data sets (TCGA, CCLE, RPE-1 and Triplosensitivity), we therefore consider these 112 genes as having strong evidence of compensation across cancer types.

In the tissue-specific analysis, we identified a total of 265 additional genes that were compensated in both CCLE and TCGA for at least one cancer type (**Fig. 2c**). This brings the total number of identified genes common to both datasets to 377, with 68 genes in both the pan-cancer and tissue-specific analysis. The additional tissue-specific genes tended to occur in one cancer type exclusively, as we observed less than 10% of compensated genes that were shared in two, and no genes that were shared in four or more tissues. Across pan-cancer and tissue-specific analyses, only some of the most compensated genes were shared between cell lines (CCLE) and tumors (TCGA), where the former showed a stronger enrichment in cell cycle and the latter a stronger enrichment in immune-related processes (**Fig. S2g-h**).

**Fig. S2.**
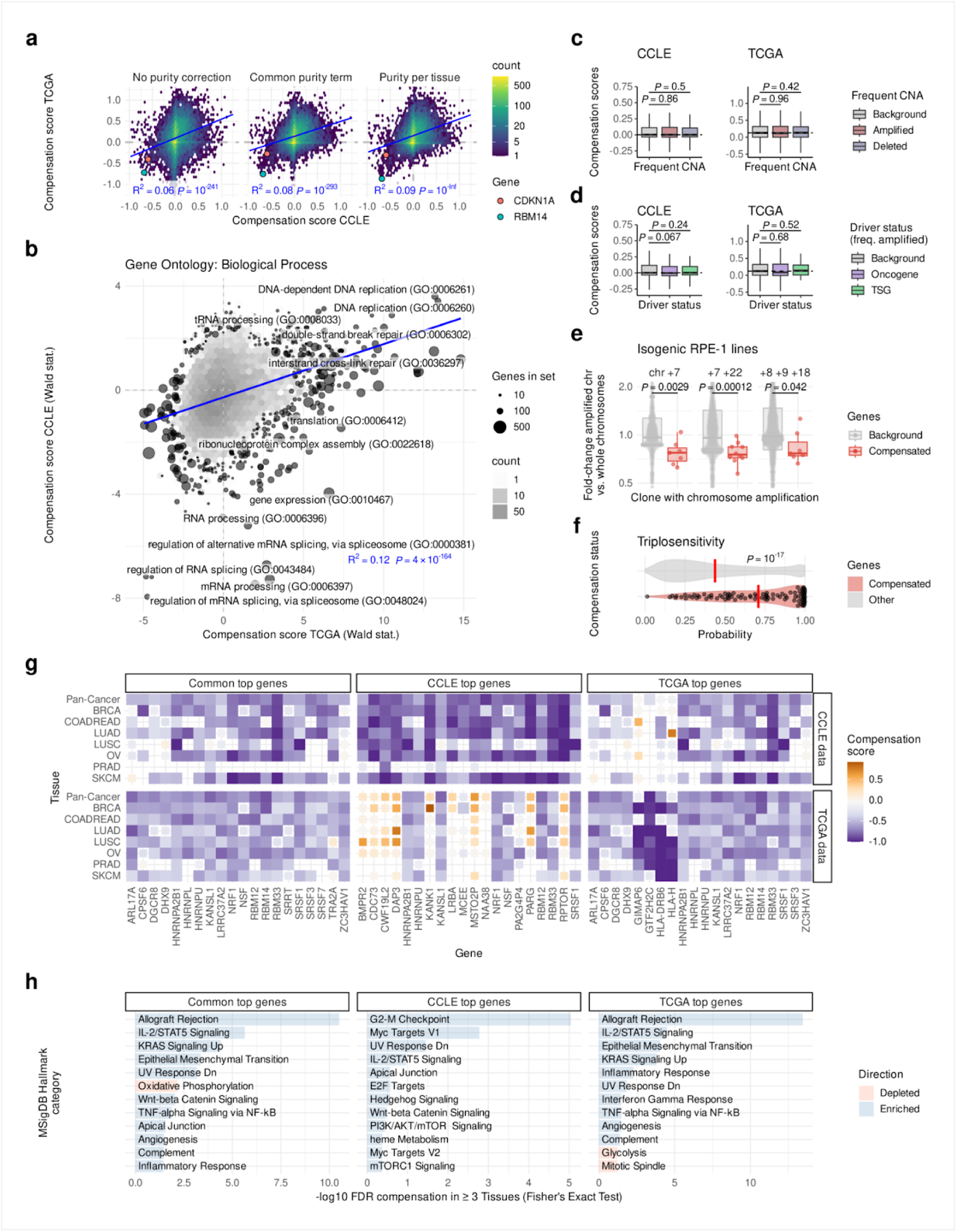
Compensation analysis. **(a)** Modeling the TCGA scaling and deviation produced the highest correlation with CCLE when modeling non-cancer cells per tissue (right). **(b)** Gene ontology enrichment of gene deregulation using a linear regression model and the Wald statistic for TCGA (x axis) and CCLE (y axis) shows commonly compensated (negative values) and hyperactivated (positive values) categories. **(c)** Frequently amplified and deleted genes, as well as **(d)** OGs and TSGs are not preferentially compensated. **(e)** Identified compensated genes are also expressed less than expected in RPE-1 clones. Fold changes were calculated over the parental RPE-1 line and then normalized per amplified chromosome. Boxes median ± quartile, whiskers 1.5x inter-quartile range. *P* -values from a linear regression model (a, b) two-sided *t* -tests (c, d) and one-sided *t* -tests (e). R^2^ Pearson correlation. **(f)** Overlap between compensated genes and genes previously identified as disease-causing when amplified (Wilcox test). **(g)** Compensation scores of common and dataset-specific genes (horizontal panels) in their respective datasets (vertical panels), as well as **(h)** enrichment of genes found in three or more analyses (Fisher’s Exact Test). Compensated genes in (g) are shown in larger squares relative to those not passing our threshold criteria.

**Fig. S3.**
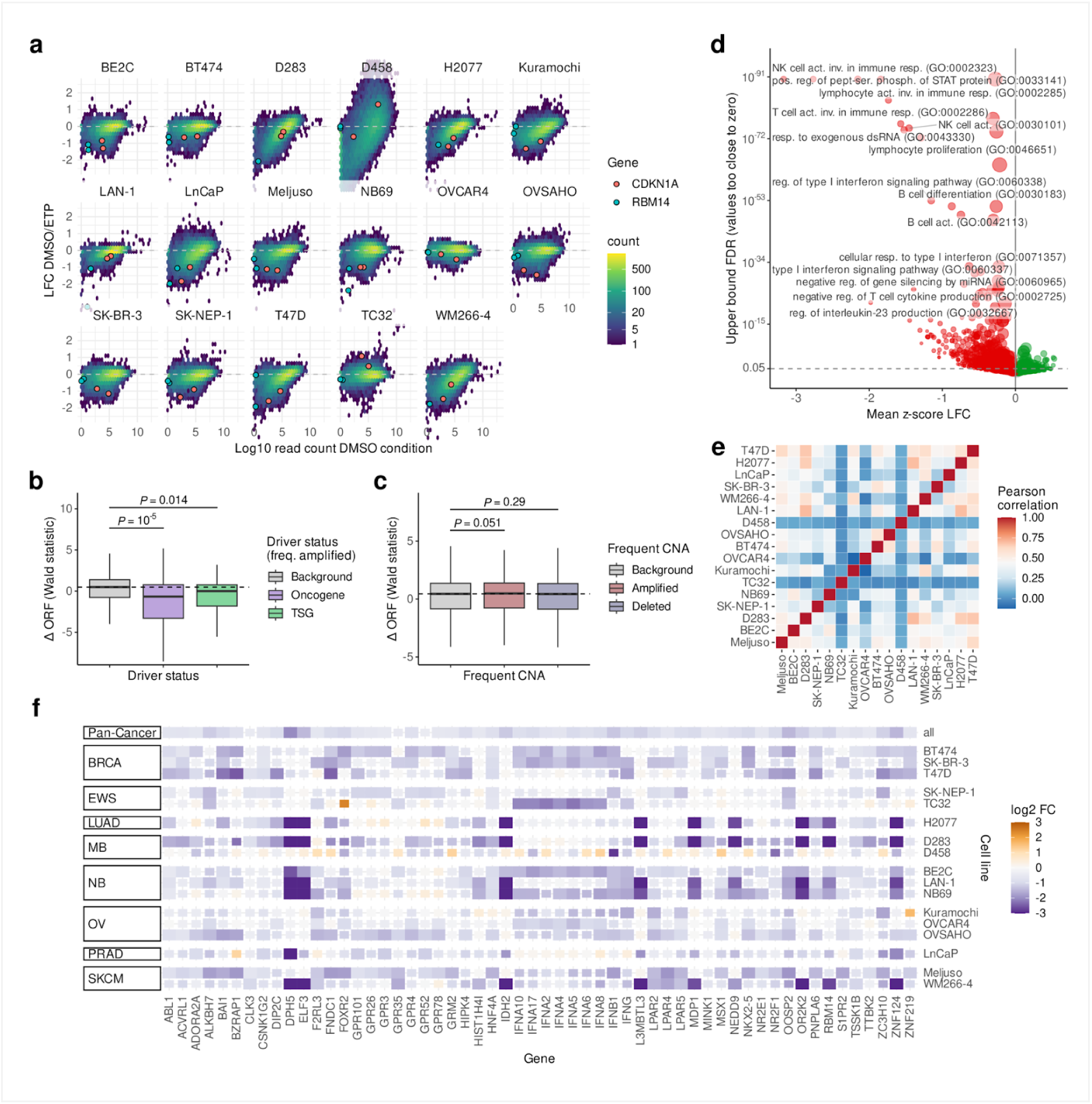
ORF screen analysis. **(a)** Overview of the expression (x axis) and dropout levels (y axis) for all screens with two genes of interest highlighted. **(b)** Both OGs and TSGs dropped out preferentially in the ORF screens over other genes. **(c)** We observed no differential ORF dropout of frequently amplified or deleted genes. **(d)** Volcano plot of Gene Ontology categories that are preferentially enriched (green, positive values) or drop out (red, negative values) across ORF screens. Boxes represent median ± quartile, whiskers 1.5x inter-quartile range. *P* -values from two-sided *t* -tests. **(e)** Pearson correlation of log fold-changes across screens shows values of 0.4 to 0.6, with three less correlated screens. We did not observe a trend where more similar tissues showed a higher correlation and hence quantified viability difference common to all screens using a linear regression model. **(f)** Gene dropout ( *log* 2 fold changes) in the pan-cancer, as well as in cell-line specific analyses. Genes common to four or more ORF screens are shown, with hits passing our thresholds indicated in larger square size.

### ORF screens reveal genes that are toxic when overexpressed

One reason a gene may be compensated is because cancer cells cannot tolerate its overexpression. To detect such situations, we aggregated gene toxicity data from 17 different ORF screens across 10 tumor types ^25–33^. In each of these ORF screens, cells were transduced with the lentiviral ORFeome library ^39^ containing 16,100 barcoded constructs that encode for a total of 12,753 genes. After transduction and construct marker selection, cells were subjected to drug treatment or vehicle control and grown for up to 3 weeks. With only the vehicle control arms, we determined the effect of each gene’s overexpression on cell viability and/or proliferation. For this, we quantified the *log2*-fold changes of lentiviral barcodes between early and late time points across all screens using pan-cancer and tissue-specific linear models (**Fig. 2d**; **Fig. S3a-b**).

Across all cancer cell lines, we observed many more genes to be depleted rather than enriched in these screens, with *DPH5*, *ZNF124*, *ABL1,* and *HNF4A* showing the most significant toxicity (**Fig. 2e**). We did not observe a preferential dropout or enrichment in commonly amplified vs. commonly deleted genes (**Fig. S3b**), but both OGs and TSGs were depleted more strongly than other genes (**Fig. S3c**). Interestingly, overexpressing OGs in a cancer cell background led to an even stronger viability defect than TSGs, in line with oncogene-induced senescence as a major driver of gene toxicity ^21,22^. However, the genes that dropped out the strongest on average were pro-inflammatory genes, indicating that their overexpression is detrimental to cancer cell growth even in *in vitro* conditions without immune cells (**Fig. S3d**).

Using these screen results, we categorized genes whose overexpression was associated with at least a 30% decrease in growth (*P* < 10^-5^) as potentially toxic. Among the 12,753 genes tested, 341 genes met these criteria across cancer types, and 442 within at least one cancer type (**Fig. 2e-f; Supp. Table 3**). The majority of the pan-cancer toxic genes (194) were also found in the tissue-specific analysis (**Fig. 2f**). To focus on the most confident hits, we considered only genes identified across the pan-cancer analysis (**Fig. S3e-f**).

### Integration of pan-cancer compensation and toxicity analyses identifies ARGOS genes

Compensated and hyperactivated genes are spread along the genome and are not enriched for frequently amplified genes. The same is true for toxic genes, *i.e.* genes that drop out in the ORF screens (**Fig. 3a**). More generally, the distribution of genes of all classes along the genome followed the overall gene density (**Fig. S4a**) and no other strong co-occurrence patterns were observed. However, compensated genes were on average also toxic when overexpressed, and hyperactivated genes promoted cell growth and survival in the ORF screens (**Fig. 3b**; **Fig. S4b-c**).

**Fig. 3.**
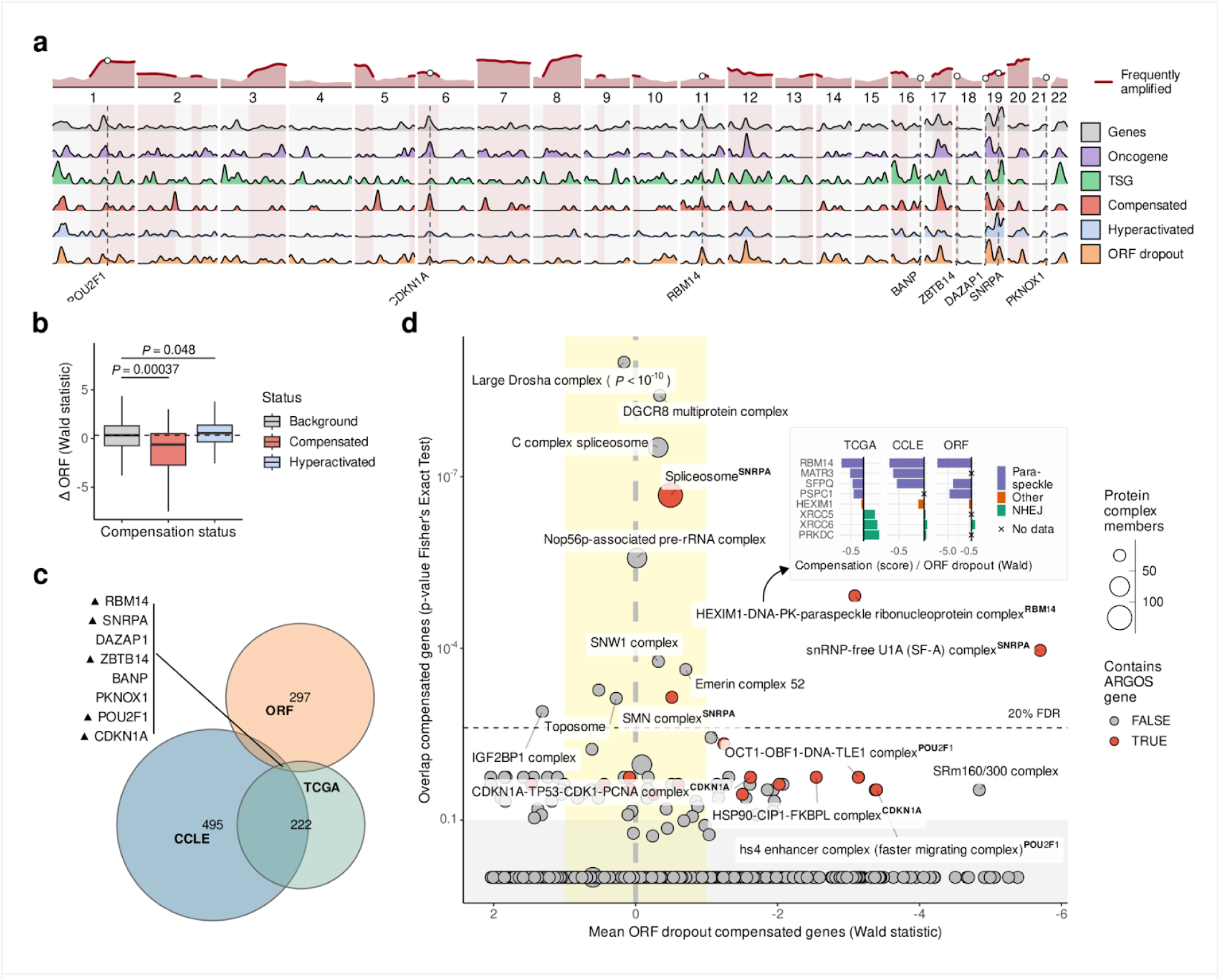
Overlap of compensation and toxicity identifies ARGOS genes. **(a)** All gene classes are spread along the genome and densities using a Gaussian estimator with a 1 Mb bandwidth show clustering only in gene-dense regions (cf. Fig. S4a). **(b)** Compensated and hyperactivated genes are respectively dropping out and increasing in ORF screens more strongly than non-regulated genes. Boxes represent median ± quartile, whiskers 1.5x inter-quartile range, *P*-values from a two-sided *t*-test. **(c)** The overlap of compensated and toxic genes identifies eight genes, five of which are amplified in over 15% of TCGA tumors (upward pointing triangles). **(d)** Prioritization of protein complexes with ARGOS genes (Fisher’s Exact test on CORUM complexes, y axis) and their toxicity (Wald statistic of a linear regression model for ORF dropout, x axis) identifies the HDP-RNP and U1A spliceosomal complexes. Paraspeckle genes in the HDP-RNP complex are both compensated and toxic, while nonhomologous end-joining (NHEJ) genes show no effect or the opposite trend.

Intersecting the sets of pan-cancer compensated and toxic genes yields eight high-confidence ARGOS genes, five of which are also frequently amplified: *RBM14*, *SNRPA*, *ZBTB14*, *POU2F1*, and *CDKN1A* (**Fig. 3a; c**). As gene dosage compensation has previously been shown to primarily occur at the level of protein complexes ^5,10,40^, we investigated their functional impact based on complex membership: Prioritizing both by the number of compensated genes within a complex and by their degree of toxicity (**Fig. 3d**), the top ‘hits’ of this analysis are *RBM14* (acting in the HEXIM1-DNA-PK-paraspeckle ribonucleoprotein complex, HDP-RNP; *FDR = 0.003*) and *SNRPA* (involved in U1A splicing; *FDR = 0.025*). By contrast, general splicing and mi/rRNA processing complexes showed compensation but no toxicity. In addition, *CDKN1A* (p21), a well-characterized TSG and a potent cell cycle inhibitor, showed strong compensation and toxicity based on our analysis, but was not significantly enriched in any specific complex. To consider both a well-known and a rarely studied gene, we chose *CDKN1A* and *RBM14* for a more detailed investigation. *CDKN1A* is part of the frequently amplified *p* arm of chromosome 6 (**Fig. 3a**), and its direct role in cell cycle arrest makes it an expected ARGOS gene that can serve as a control for our identification and validation strategies. *RBM14* is located at the edge of a focal amplification next to *CCND1* on chromosome 11 and the nature of its toxicity is unknown. However, its complex members suggest that it may be implicated in the cellular DNA damage response and in the activation of innate immune signaling (**Fig. 3d**).

**Fig. S4.**
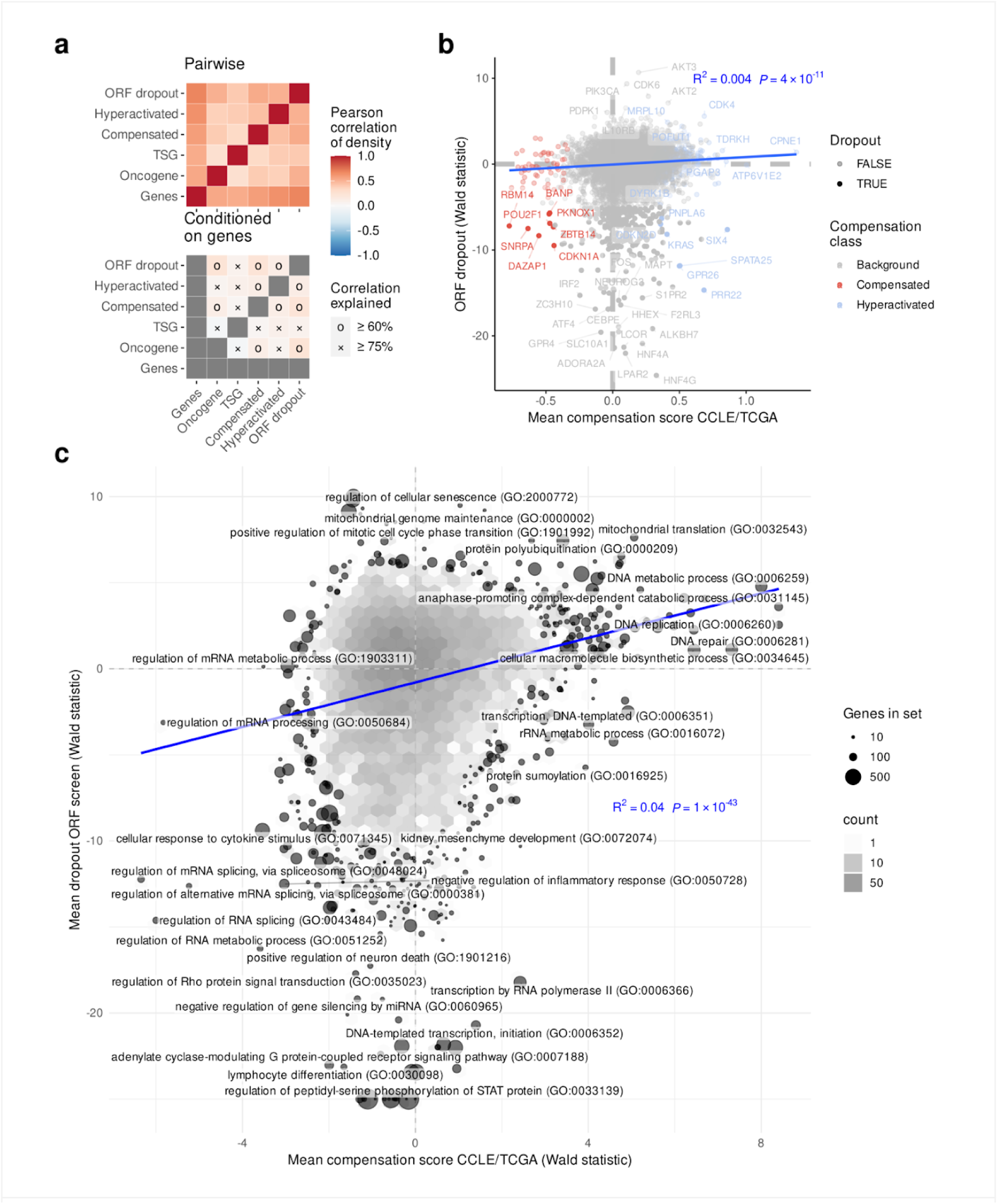
Characterization of compensation and ORF overlap. **(a)** Pearson correlation of the gene densities in Fig. 3a (top) and adjusted for gene densities (bottom) shows that gene density along the genome explains most of the observed correlations. **(b)** Compensation vs. ORF dropout plot with genes in the respective compensation categories highlighted. **(c)** Correlation of Gene Ontology categories for compensation scores (x axis) and toxicity (y axis; limited at -25). R^2^ Pearson correlation, P-values from linear regression models.

### The well-known cell cycle inhibitor *CDKN1A* as an ARGOS gene

Amplification of *CDKN1A* occurs at the arm-level in chromosome 6p. Copy number gains of known oncogenes that may drive this arm-level gain ^41^, such as *CCND3*, *POU5F1*, and *PIM1* reside in its close vicinity (**Fig. 4a**). Our analysis identified this tumor suppressor gene as a compensated ARGOS gene in TCGA and CCLE (**Fig. 4b**), and our ORF screen data pointed to overexpression of *CDKN1A* resulting in a toxic effect (**Fig. 4b-c**), in agreement with its important role in cell cycle arrest. To functionally validate the toxic effect of *CDKN1A* overexpression, we selected cancer cell lines from two tumor types (lung and breast): one with normal copy number and average mRNA expression (“non-compensated”) and one with lower mRNA/protein expression levels than expected by its amplification (“compensated”) (**Fig. 4d**; **Fig. S5a**). We transduced SK-LU-1 lung adenocarcinoma and MDA-MB-231 breast cancer cells with doxycycline-inducible and V5-tagged constructs encoding either *CDKN1A* or a luciferase control. By treating cells with a range of doxycycline concentrations (50-500 ng/mL), we were able to detect increases in *CDKN1A* transcript and protein levels (up to 3-fold) for both cell lines in a dose-dependent manner (**Fig. S5b-c**). A luciferase reporter assay confirmed luciferase expression in each cell line model following doxycycline-mediated induction (**Fig. S5d**). To validate the deleterious impact of *CDKN1A* overexpression on cell proliferation, we performed a live-cell imaging assay where we monitored cell confluency in *CDKN1A-* or luciferase-overexpressing cells over time. Indeed, increased levels of gene expression led to a decrease in cell proliferation, with >60% growth inhibition observed for SK-LU-1^CDKN1A^ and MDA-MB-231^CDKN1A^ cells treated with 500 ng/mL doxycycline (**Fig. 4e**), independent of amplification status. These results are in line with the well-established tumor-suppressive role of CDKN1A ^42^. Taken together, these findings serve as proof of concept that we can generate cellular models of gene overexpression to validate ARGOS candidates identified by our compensation and toxicity analyses.

**Fig 4.**
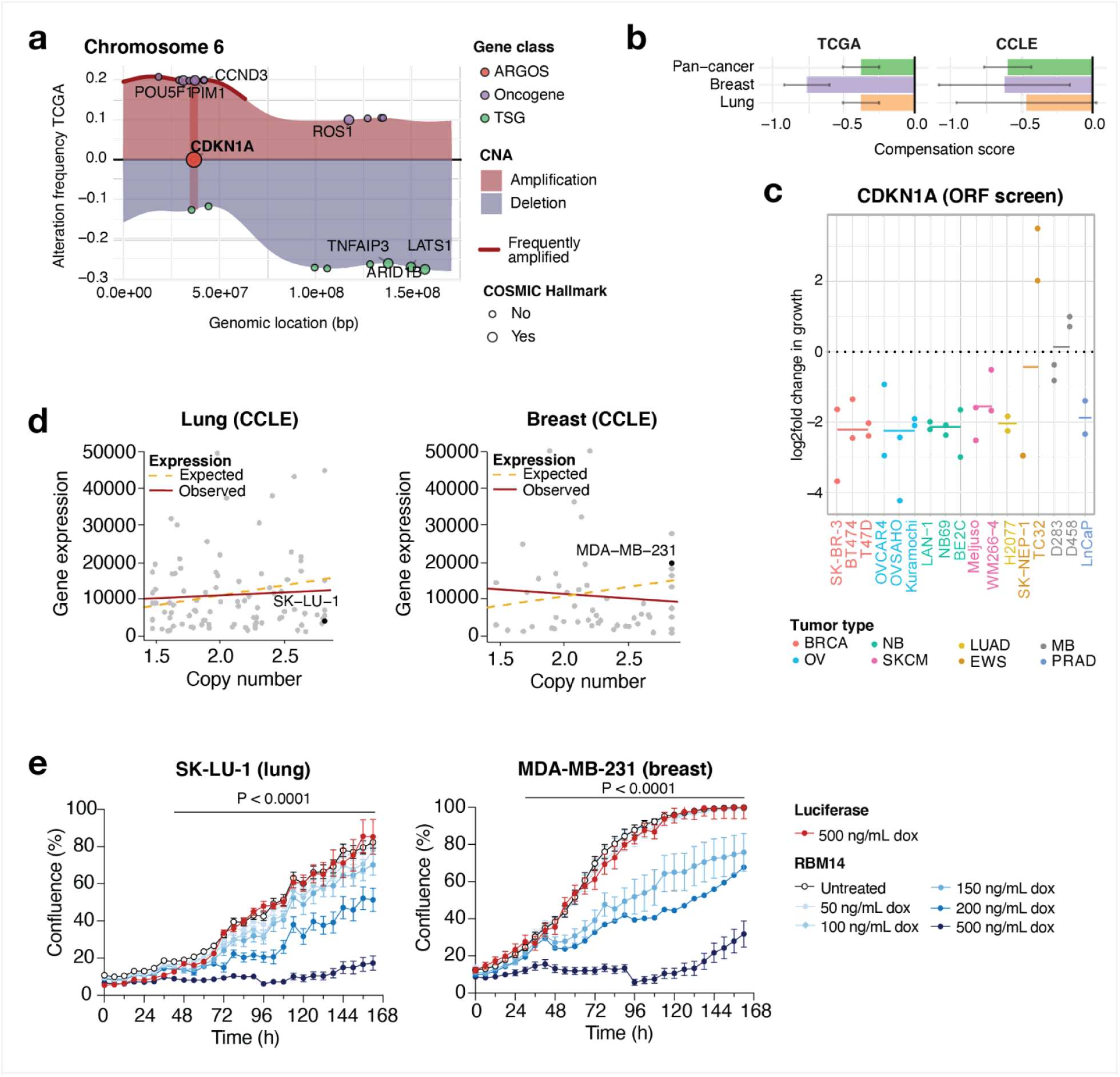
CDKN1A as an ARGOS gene. **(a)** The map of copy number alterations in chromosome 6 shows a p-arm amplification, where *CDKN1A* is located among other known oncogenes. Amplifications (red) and deletions (blue) are shown for this genomic region. **(b)** Compensation scores for *CDKN1A* in our pan-cancer analysis (green) as well as in breast (purple) and lung (orange) lineages chosen for downstream functional validation. Bars represent the mean, error bars the standard deviation of the posterior. **(c)** Depletion scores for *CDKN1A* ORFs across 17 independent screens, including breast (BRCA), ovarian (OV), neuroblastoma (NB), skin cutaneous melanoma (SKCM), lung adenocarcinoma (LUAD), Ewing sarcoma (EWS), medulloblastoma (MB) and prostate adenocarcinoma (PRAD) cell lines. The mean log-fold change in growth is shown as a line for each tumor type. **(d)** Lung and breast cancer cell lines chosen for functional validation based on their gene expression and DNA copy number profile on CCLE. **(e)** CDKN1A overexpression leads to a growth inhibition phenotype upon varying levels of overexpression. The mean and S.D. of three replicates are shown. Data analyzed using two-way ANOVA.

**Fig S5.**
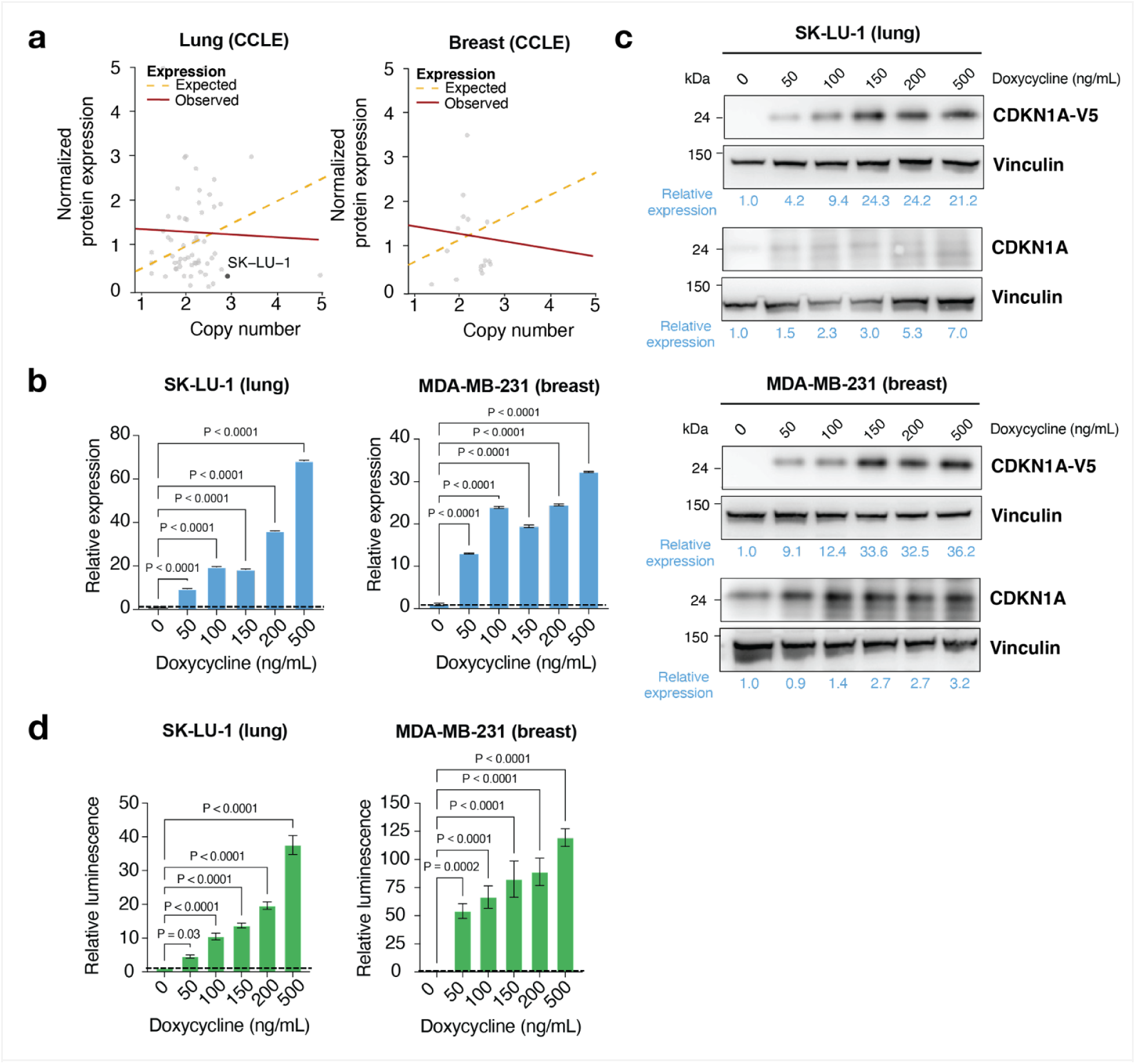
Generation of doxycycline-inducible cell model systems to study CDKN1A overexpression. **(a)** CDKN1A protein expression level and copy number for lung and breast cancer cell lines in CCLE. No data for MDA-MB-231 cells available. **(b)** Increasing levels of *CDKN1A* transcript were detected by RT-qPCR in SK-LU-1^CDKN1A^ and MDA-MB-231^CDKN1A^ cells when treated with increasing concentrations of doxycycline. Expression values are shown relative to untreated (0 ng/mL) control. Mean and standard deviation of three replicates. Data analyzed with one-way ANOVA. **(c)** Increases in exogenous (V5-tagged) and total CDKN1A protein were detected by immunoblotting in the same cell models. Vinculin was used as loading control. Expression values are shown relative to untreated (0 ng/mL) control. **(d)** Activity of luciferase cells was determined by a reporter assay. Increases in luminescence were detected for both cell lines at increasing concentrations of doxycycline. Luminescence values are shown relative to untreated (0 ng/mL) control. Mean and standard deviation of three replicates. Data analyzed with one-way ANOVA.

### RBM14 overexpression reduces proliferation of human lung and breast cancer cell lines and causes cell death by apoptosis

RBM14 belongs to the family of RNA binding proteins, which interact with RNA transcripts to regulate their splicing, cytosolic transport and translation ^43,44^. RBM14 plays a role in alternative splicing, regulation of DNA repair via the canonical non-homologous end joining (c-NHEJ) pathway, and is part of the HEXIM1-DNA-PK-paraspeckle components-ribonucleoprotein (HDP-RNP) complex that regulates innate immune response through cGAS-STING signaling ^45–47^. Its depletion leads to disrupted genome integrity and the accumulation of DNA damage during mouse embryogenesis ^48^, and hinders mitotic spindle assembly and chromosome segregation in human U2OS bone osteosarcoma cells ^49^.

RBM14 amplification occurs as a focal event in chromosome 11, close to copy number gains of the oncogene *CCND1* (**Fig. 5a)**. Our analysis showed strong gene compensation in breast and lung cancer (**Fig. 5b**), and ORF screens pointed to RBM14 overexpression also leading to detrimental effects on cell proliferation in multiple cancer types (**Fig. 5c**). To validate the cellular effects of *RBM14* overexpression in human cell lines, we selected two lung adenocarcinoma (NCI-H838; NCI-H1650) and two breast cancer (ZR-75-1; HCC70) cell lines that either exhibit high copy number and lower-than-expected mRNA/protein expression levels (“compensated”) or remain euploid at the *RBM14* locus while expressing average levels of gene transcript and protein (**Fig. 5d**; **Fig S3a**). To generate cell models of RBM14 overexpression, we first transduced cells with a doxycycline (dox) inducible V5-tagged construct encoding either RBM14 protein (generating NCI-H838^RBM14^, NCI-H1650^RBM14^, ZR-75-1^RBM14^, and HCC70^RBM14^) or a luciferase overexpression control (generating NCI-H838^luc^, NCI-H1650^luc^, ZR-75-1^luc^, and HCC70^luc^). By treating all four RBM14-overexpressing cell lines with increasing concentrations of dox (0 to 500 ng/mL), we were able to detect a dose-dependent induction in *RBM14* gene transcript as well as increased expression of V5-tagged exogenous and total RBM14 protein (**Fig. S6b-c**). We confirmed luciferase activity in control cell lines by measuring luminescent signal in the presence of substrate following doxycycline treatment (**Fig. S6d**).

**Fig 5.**
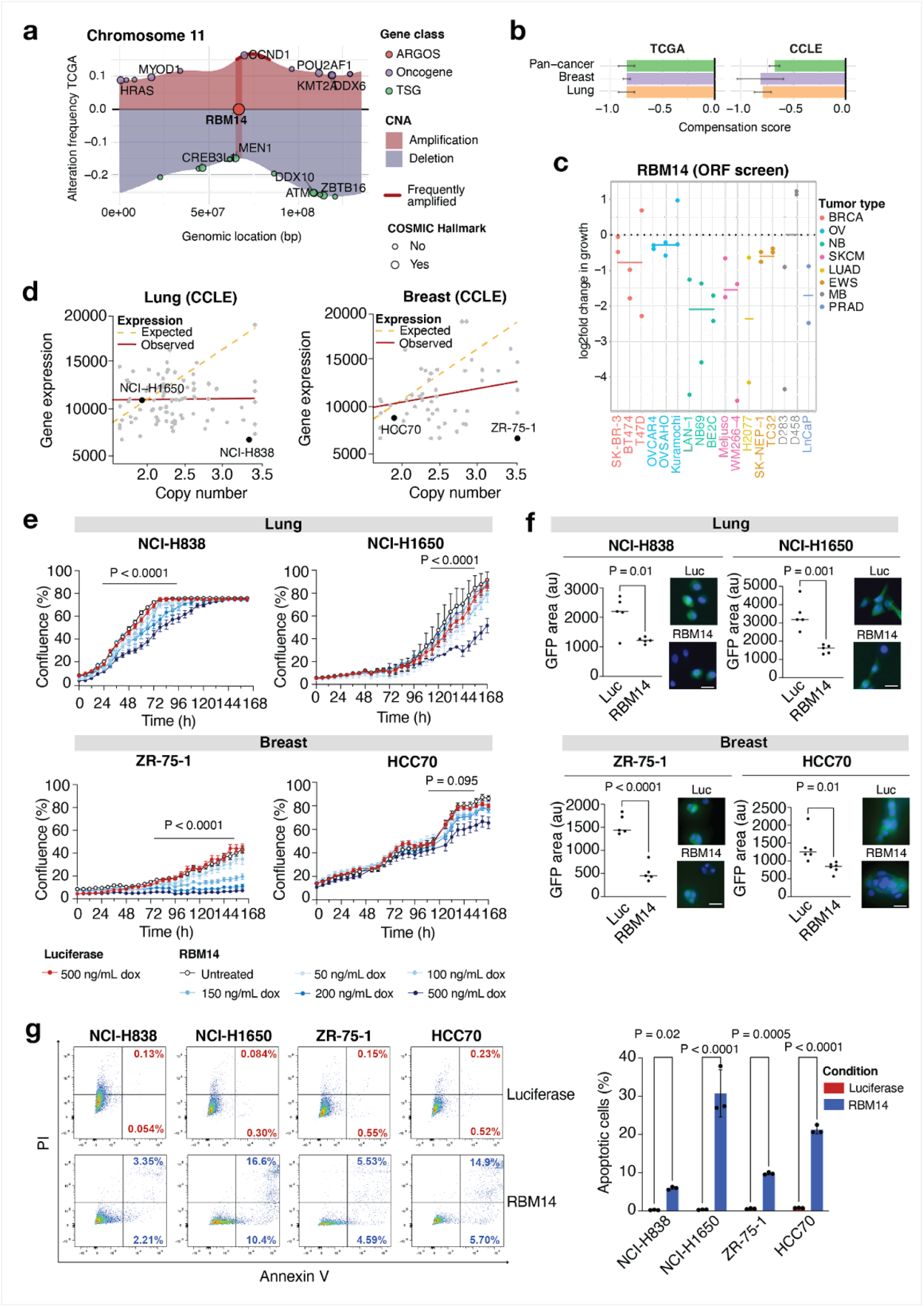
RBM14 as an ARGOS gene. **(a)** The map of copy number alterations in chromosome 11 shows a focal amplification, where *RBM14* is located next to *CCND1*. Amplifications (red) and deletions (blue) are shown for this genomic region. **(b)** Compensation or ORF dropout scores for *RBM14* in our pan-cancer analysis (green) as well as in breast (purple) and lung (orange) lineages chosen for downstream functional validation. Bars represent the mean, error bars the standard deviation of the posterior. **(c)** Depletion scores for RBM14 ORFs across 17 independent screens, including breast (BRCA), ovarian (OV), neuroblastoma (NB), skin cutaneous melanoma (SKCM), lung adenocarcinoma (LUAD), Ewing sarcoma (EWS), medulloblastoma (MB) and prostate adenocarcinoma (PRAD) cell lines. The mean log-fold change in growth is shown as a line for each tumor type. **(d)** Lung and breast cancer cell lines chosen for functional validation based on their gene expression and DNA copy number profile on CCLE. **(e)** RBM14 overexpression leads to a growth inhibition phenotype. The mean and S.D. of three replicates are shown. Data analyzed using two-way ANOVA. **(f)** RBM14 overexpressing lung and breast cells show a decrease in nascent protein synthesis. Fluorescence (GFP; green) is quantified relative to cell number (DAPI; blue). Luciferase overexpressing cells were used as control. Scale bar: 100 μ M. Data analyzed by unpaired t-test. **(g)** The fraction of apoptotic cells following RBM14 overexpression was determined by flow cytometry-based quantification of annexin V and PI positive cells. The mean and S.D. of three replicates is shown. Data analyzed by unpaired t-test.

We next evaluated the impact of *RBM14* overexpression on cell proliferation using live-cell imaging of the RBM14- and luciferase-overexpressing cell lines treated with increasing concentrations of dox. All RBM14-overexpressing cell lines exhibited a dose-dependent reduction in proliferation, with the strongest effects observed as early as 48 h following 500 ng/mL dox induction. By seven days of treatment, NCI-H838^RBM14^, NCI-H1650^RBM14^, ZR-75-1^RBM14^, and HCC70^RBM14^ cells showed >50% decrease in confluency (*P* < 0.0001) when compared to their respective luciferase controls, supporting the observation that *RBM14* overexpression has inhibitory effects on cell proliferation *in vitro* (**Fig. 5e**). This growth inhibitory phenotype was supported by the observation that RBM14 overexpression interfered with nascent protein synthesis, which we measured by incubating cells with a fluorescently labeled methionine analog. Following 48 h of dox induction, we performed immunofluorescence and quantified the amount of average fluorescent signal detected per cell (**Fig. 5f**). We found that RBM14 overexpression caused a two- to three-fold decrease in fluorescent signal, and hence nascent protein synthesis, when compared to luciferase controls (*P* = 0.0106 for NCI-H838; *P* = 0.0011 for NCI-H1650; *P* < 0.0001 for ZR-75-1; *P* = 0.0112 for HCC70). We next explored whether the reduction in cell proliferation associated with RBM14 overexpression is a result of cell cycle arrest or cell death. Flow cytometry for BRdU incorporation did not indicate significant changes in cell cycle profile between RBM14- or luciferase- overexpressing NCI-H838, NCI-H1650, ZR-75-1, and HCC70 cells (**Fig. S7a-b**). However, when we performed flow cytometry for Annexin V/PI, 20-30% of cells were positive for Annexin V in NCI-H1650^RBM14^ and HCC70^RBM14^ cells by 72 h post induction with 500 ng/mL doxycycline (*P* < 0.0001) while apoptotic rates remained < 3% in luciferase controls (**Fig. 5g**). The apoptosis rates of NCI-H838^RBM14^ and ZR-75-1^RBM14^ were lower, with 5-10% cells being positive for Annexin V, but these were still significantly higher rates of apoptosis in comparison to their luciferase controls (*P* = 0.025 for NCI-H838; *P* = 0.0005 for ZR-75-1). Altogether, these results show that low-level overexpression of RBM14 negatively impacts cellular proliferation and viability in our lung and breast cancer models, regardless of gene amplification status.

**Fig S6.**
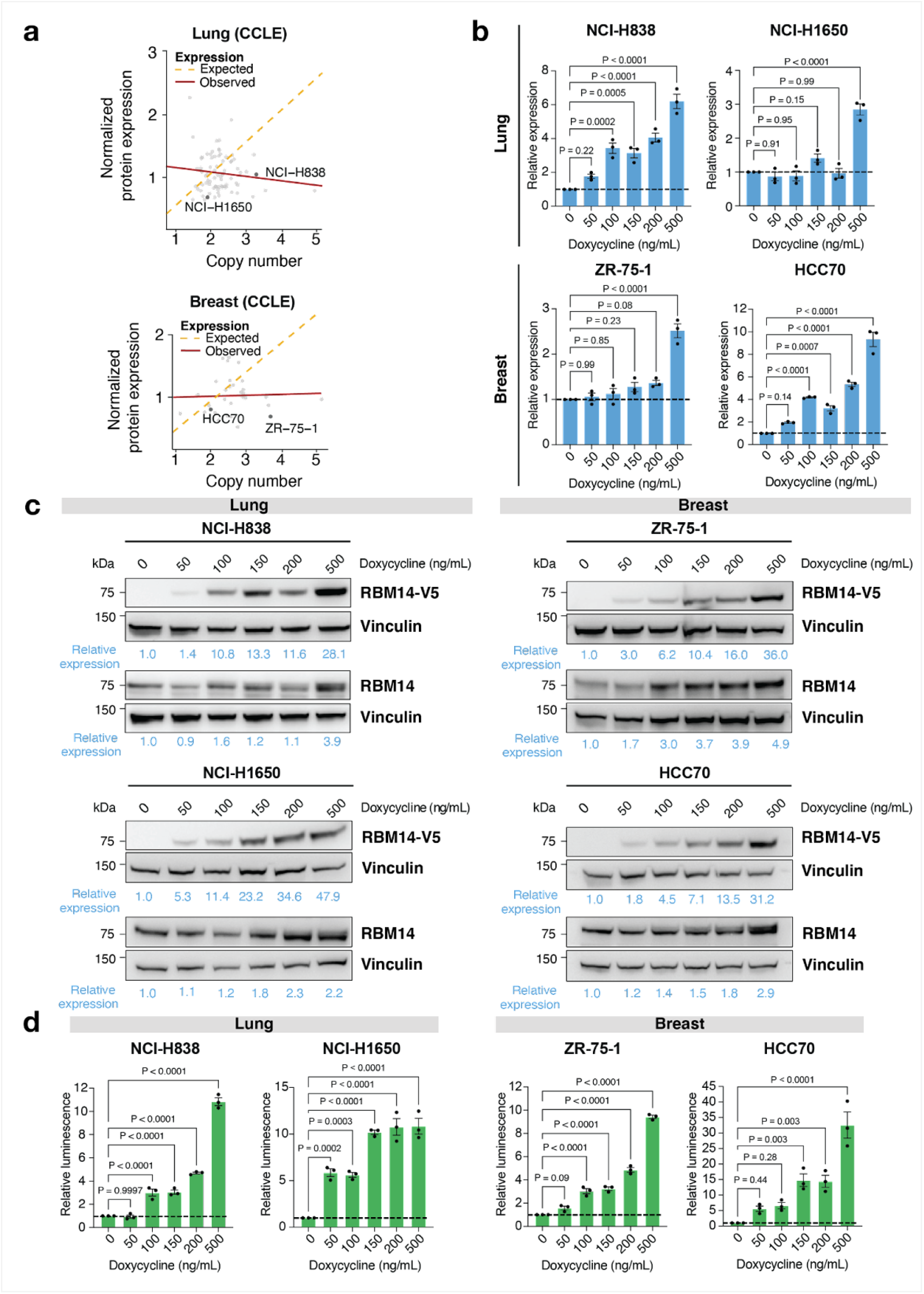
Generation of doxycycline-inducible cell model systems to study RBM14 overexpression. **(a)** Cell lines chosen for functional validation show similar compensation profiles based on protein expression level and copy number. **(b)** Increasing levels of *RBM14* transcript were detected by RT-qPCR in NCI-H838^RBM14^, NCI-H1650^RBM14^, ZR-75-1^RBM14^ and HCC70 RBM14 cell lines when treated with increasing concentrations of doxycycline. Expression values are shown relative to untreated (0 ng/mL) control. Mean and standard deviation of three replicates. Data analyzed with one-way ANOVA. **(c)** Increases in exogenous (V5-tagged) and total RBM14 protein were detected by immunoblotting in all four cell models. Vinculin was used as loading control. Expression values are shown relative to untreated (0 ng/mL) control. **(d)** Activity of luciferase cells was determined by a reporter assay. Increases in luminescence were detected for both cell lines at increasing concentrations of doxycycline. Luminescence values are shown relative to untreated (0 ng/mL) control. Mean and standard deviation of three replicates. Data analyzed with one-way ANOVA.

### RBM14 overexpression increases reliance on DNA repair by c-NHEJ and activates cGAS-STING signaling

Our analysis of compensated protein complexes identified RBM14 as one of several paraspeckle proteins interacting with the DNA-dependent protein kinase (DNA-PK), and other members of the HDP-RNP complex (cf. **Fig. 3d**). This subnuclear complex mediates DNA damage sensing and repair and regulates cGAS-STING signaling and the innate immune response ^47,50,51^. We therefore explored whether DNA damage is a mechanism by which RBM14 overexpression might lead to cell death. We first exposed the lung adenocarcinoma NCI-H838^RBM14^/NCI-H838^luc^ and NCI-H1650^RBM14^/NCI-H1650^luc^ cell line pairs to 2 Gy ionizing radiation (IR), and by immunofluorescence measured levels of nuclear γ-H2AX foci as a biomarker of DNA damage. For both cell lines, we were able to detect an initial increase in the number of γ-H2AX foci 15 min after IR, followed by a gradual decrease over time and a return to baseline levels by 360 min after exposure (**Fig. 6a**). Interestingly, NCI-H838^RBM14^ and NCI-H1650^RBM14^ cells showed a 30-50% reduction of γ-H2AX foci when compared to their respective luciferase controls, with the strongest effect at 60 min post-IR (*P* < 0.0001). We observed a similar decrease in ZR-75-1^RBM14^/ZR-75-1^luc^ and HCC70^RBM14^/HCC70^luc^ pairs of breast cancer cell lines, albeit with a lower effect size (**Fig. S8a**).

**Fig 6.**
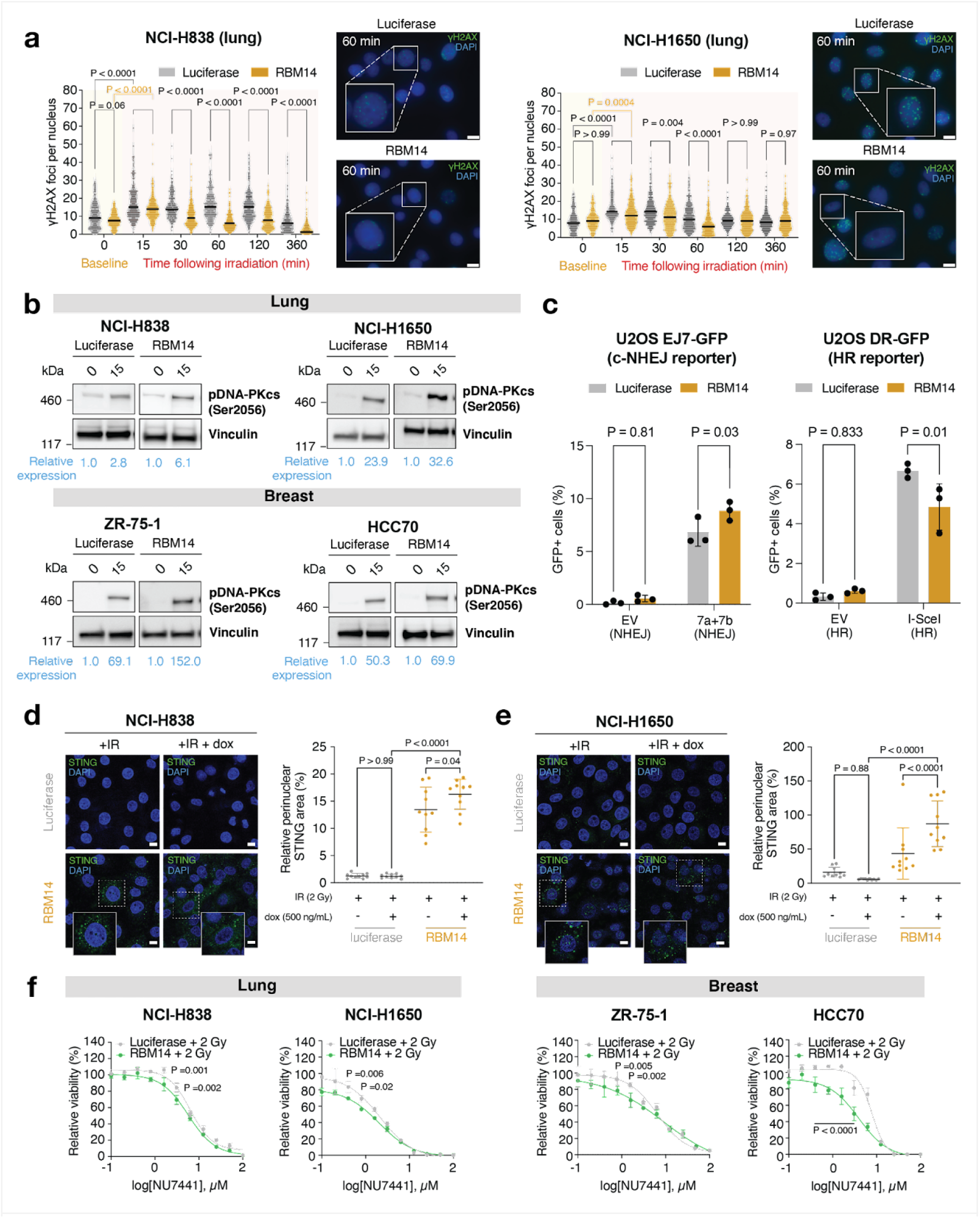
RBM14 overexpression leads to differential DNA damage response and STING activation. **(a)** The number of yH2AX foci was quantified by immunofluorescence in lung NCI-H838^RBM14/luc^ and NCI-H1650^RBM14/luc^ cells following 2 Gy ionizing radiation (IR). Data analyzed by two-way ANOVA. Representative images are shown for each condition at 60 min following IR. Scale bar: 10 µM. **(b)** Levels of DNA-PKcs (pSer^2056^) and RAD51 protein were detected and quantified by immunoblotting in cells at 15 min and 6h following 2 Gy IR. **(c)** RBM14 overexpressing U2OS cells exhibit higher rates of c-NHEJ-mediated repair. The fraction of U2OS^RBM14/luc^ cells that repair double-strand breaks by c-NHEJ (EJ7-GFP; left) or HR (DR-GFP; right) was determined by quantifying GFP-positive cells in a flow cytometry-based reporter assay. GFP-positive cells were normalized to GFP transfection controls. Empty vector (EV) controls were included for comparison. Data analyzed with unpaired t-test. **(d-e)** Immunofluorescence quantification of perinuclear STING area in RBM14- and luciferase-overexpressing **(d)** NCI-H838 and **(e)** NCI-H1650 cells. STING area was calculated relative to total nuclei area in a total of 10 images per condition. Data analyzed with two-way ANOVA. Scale bar: 10 μ M. **(f)** Response of lung and breast cell line models to DNA-PK inhibition. Cells were irradiated with 2 Gy and treated with the small-molecule NU7441 for 72h. Cell viability relative to DMSO control was assessed in a CellTiterGlo luminescence assay. Data analyzed with two-way ANOVA.

Given RBM14’s interaction with other c-NHEJ protein members in our analysis and its reported involvement in this DNA repair pathway ^46^, we hypothesized that the difference we observed in γ-H2AX phosphorylation dynamics could reflect an increased reliance of RBM14-overexpressing cells on c-NHEJ over homologous recombination (HR) for resolving DNA damage. Indeed, repair by c-NHEJ occurs more rapidly than HR and can take place throughout the cell cycle as it does not require a template ^52,53^. To assess this hypothesis, we first irradiated cells with 2 Gy for 15 min and quantified changes in expression of the catalytic subunit of the DNA-PK complex (DNA-PKcs), as auto-phosphorylation at the Ser2056 (pSer^2056^) residue is known to specifically enable c-NHEJ ^54^. Following IR, we found an average of 1.8-fold higher levels of pSer^2056^ DNA-PKcs in RBM14-overexpressing cells when compared to their respective luciferase controls (**Fig. 6b**). Our second experiment leveraged the EJ7-GFP ^55^ and DR-GFP ^56^ reporter systems engineered in U2OS cells to quantify the fraction of cells that repair double-stranded breaks (DSBs) by c-NHEJ (based on CRISPR-Cas9 cutting with ‘7a’ and ‘7b’ RNA guides) and HR (based on cutting with the endonuclease I-SceI), respectively. Following transduction of U2OS cells with RBM14 or luciferase inducible vectors, we transfected cells with plasmids encoding GFP, empty vector controls for both systems, and ‘7a’ and ‘7b’ CRISPR guides (c-NHEJ assay) or I-SceI (HR assay). After 72h, we detected a significant increase in the fraction of GFP-positive U2OS EJ7-GFP RBM14-overexpressing cells repairing DSBs by c-NHEJ (*P* = 0.0280) and a significant decrease in HR-mediated repair in U2OS DR-GFP cells (*P* = 0.0142) compared to luciferase control. We did not detect changes in GFP expression for cells transfected with empty vector controls (**Fig. 6c**; **Fig. S8b**).

As c-NHEJ is an error-prone DNA repair pathway ^52^, we hypothesized that RBM14 overexpression would lead to increased misrepair following DNA damage, with resulting genomic instability. To confirm this, we first treated RBM14- and luciferase-overexpressing cells with 2 Gy IR. Next, we used time-lapse microscopy to monitor individual cells during their course of division. Upon induction of DNA damage, we observed a higher percentage of cells with mitotic defects (*P* = 0.039) in RBM14-overexpressing cells compared to luciferase controls. The mitotic aberrations detected in RBM14-overexpressing cell lines included the formation of DNA bridges during metaphase, an increase in number of micronuclei, and incomplete cellular divisions lacking metaphase alignment (**Fig. S8c-d**). These results indicate that RBM14 overexpression leads to a higher rate of aberrant mitotic division following DNA damage and suggest a mechanism by which RBM14 overexpression ultimately leads to cell death in our cell line models.

As part of the HDP-RNP complex, RBM14 also plays an important role in recognizing the accumulation of cytoplasmic DNA and is required for the production of type I interferons following cGAS-STING pathway activation ^47,57,58^. We therefore hypothesized that, in the presence of DNA damage, RBM14 overexpression would also lead to increased recognition of cytosolic DNA strands and activation of cGAS-STING signaling. To assess this, we exposed NCI-H838^RBM14^/NCI-H838^luc^ and NCI-H1650^RBM14^/NCI-H1650^luc^ overexpressing cells to 2 Gy IR and measured by immunofluorescence the area of STING outside the cellular nucleus as a marker of pathway activation ^59^. In both cell lines, RBM14 overexpression led to a significant increase in perinuclear localization of STING (quantified as relative STING area outside the cellular nucleus; *P* = 0.045 for NCI-H838^RBM14^ and *P* < 0.0001 for NCI-H1650^RBM14^), whereas no effect was observed in luciferase-overexpressing control cells (**Fig. 6d, e**). In sum, our results show that RBM14 overexpression influences DNA repair dynamics and cGAS-STING signaling in the context of DNA damage ^47^.

Finally, we explored whether the increased dependency of RBM14-overexpressing cells on c-NHEJ-mediated DNA repair exposed new vulnerabilities in these cells. For this purpose, we tested the sensitivity of our cell line models to the DNA-PK inhibitor NU7441 (thereby inhibiting c-NHEJ repair) and the RAD51 inhibitor B02 (inhibiting HR-mediated repair) after inducing DNA damage by 2 Gy IR. All four RBM14-overexpressing cell lines showed increased sensitivity to DNA-PK inhibition when compared to luciferase controls after 3 days of treatment, albeit with a mild effect size (**Fig. 6f**). By contrast, RBM14 overexpression did not alter the sensitivity to RAD51 inhibition in three out of the four cell lines (**Fig. S8e**). Altogether, these results support an increased dependency of RBM14-overexpressing cells on c-NHEJ over homologous recombination (HR) for resolving DNA damage, which underlies RBM14 toxicity, and may suggest therapeutic sensitivities associated with its expression.

**Fig S7.**
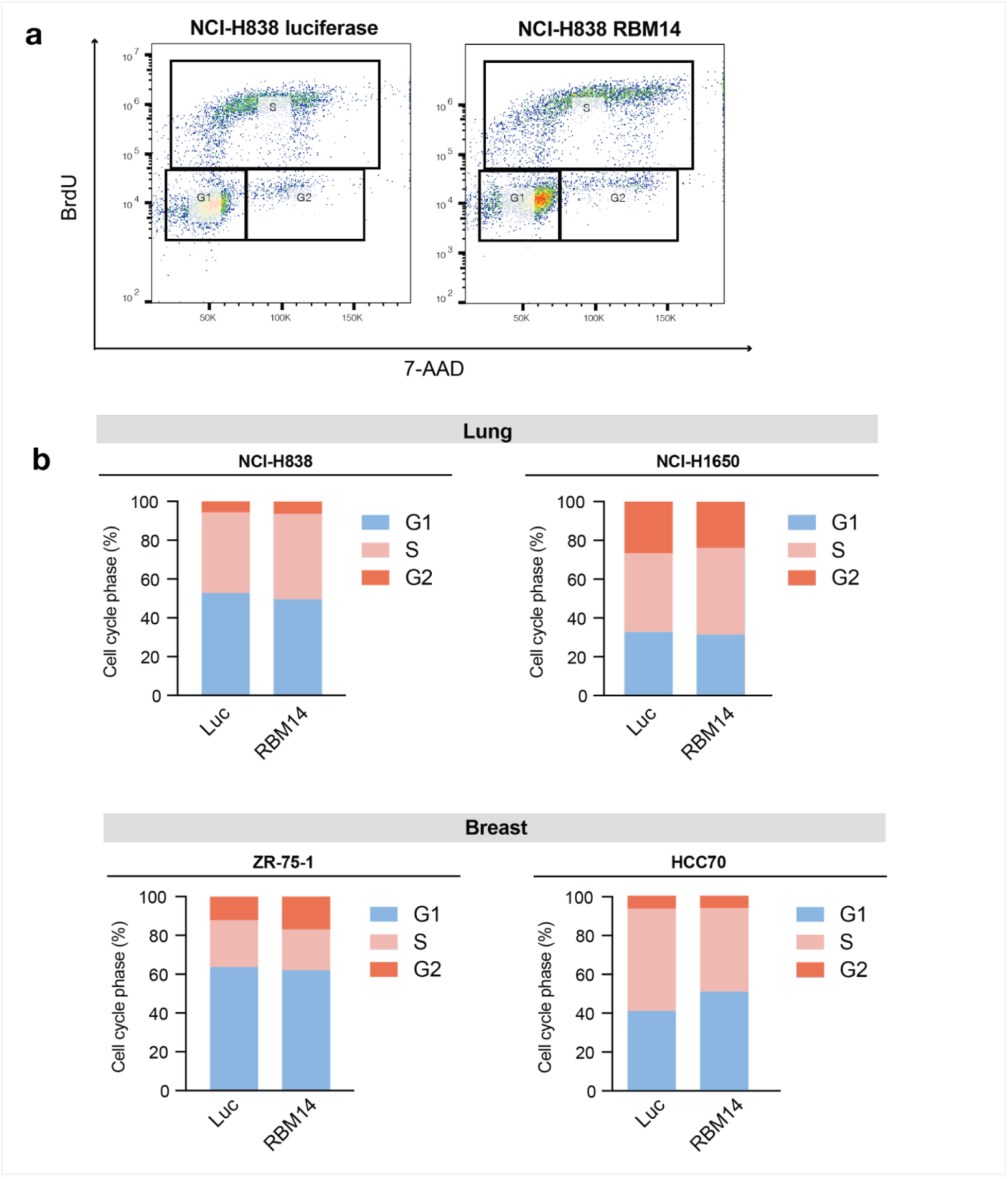
Cell cycle analysis in compensated cell line models of RBM14 overexpression. **(a)** Representative image of a BrdU incorporation assay in NCI-H838 cells overexpressing RBM14 or luciferase control. In this assay, 7-AAD was used as nucleic acid staining to quantify total DNA content. **(b)** Percentage of cells in each phase of the cell cycle following RBM14 or luciferase overexpression in lung (NCI-H838, NCI-H1650) and breast (ZR-75-1, HCC70) cancer cell lines.

**Fig S8.**
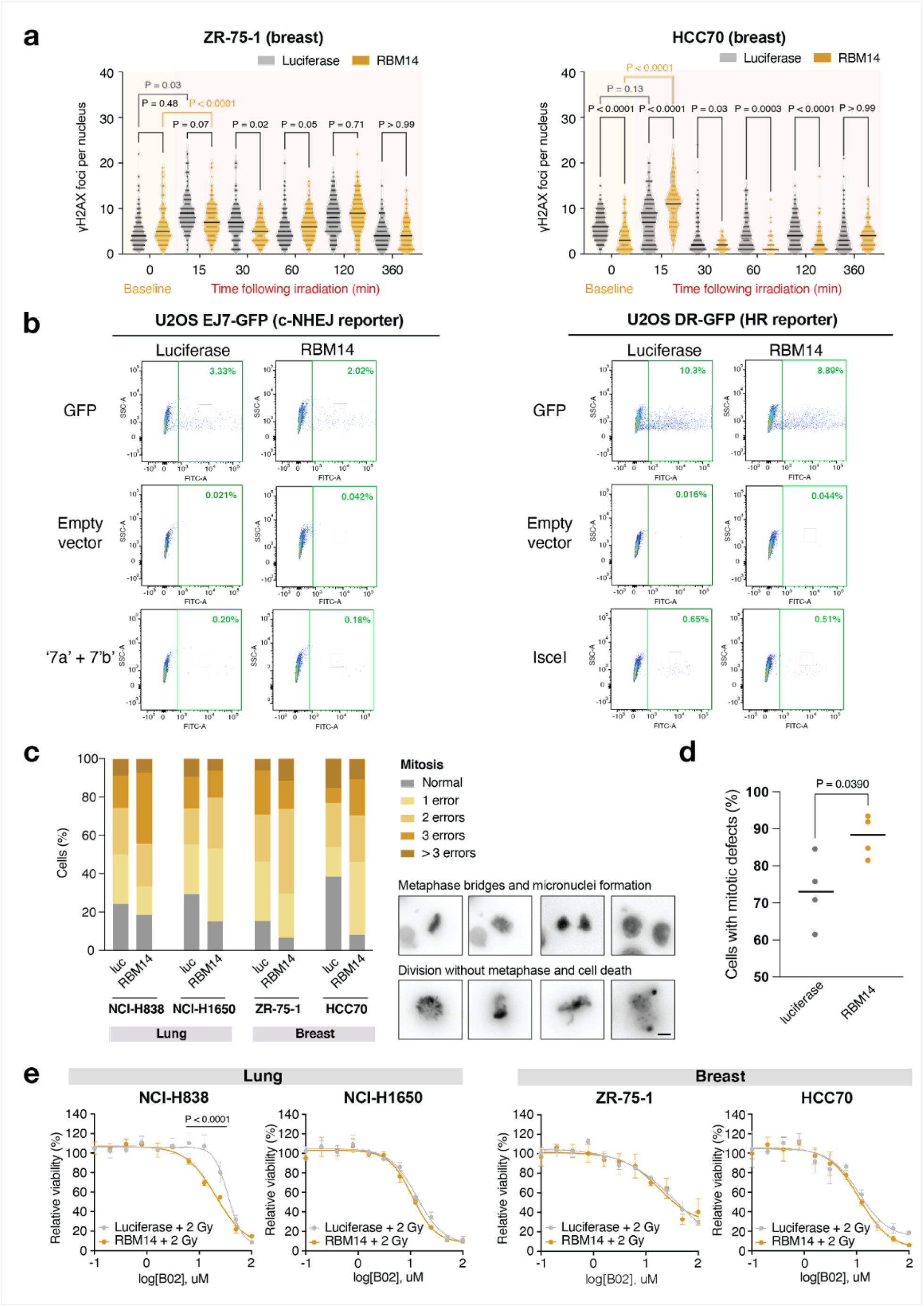
DNA damage response in cell line models of RBM14 overexpression. **(a)** The number of yH2AX foci was quantified by immunofluorescence in breast ZR-75-1^RBM14/luc^ and HCC70^RBM14/luc^ cells following 2 Gy ionizing radiation (IR). Data analyzed by two-way ANOVA. **(b)** Representative images for each condition of the DNA damage reporter assay described in Fig. 6c. The fraction of GFP-positive cells are shown in green rectangles. **(c)** Quantification of the number of mitotic defects per cell after 2 Gy IR. Representative images of observed mitotic alterations, including DNA bridges, micronuclei formation and cell division without metaphase (images on the right panel). Scale bar: 12 μM. **(d)** Rates of cells with mitotic defects following DNA damage by 2 Gy IR. The mean of each condition is shown in black. Data analyzed with unpaired t-test.. **(e)** Response of lung and breast cell line models to RAD51 inhibition. Cells were irradiated with 2 Gy and treated with the small-molecule B02 for 72h. Cell viability relative to DMSO control was assessed in a CellTiterGlo luminescence assay. Data analyzed with two-way ANOVA.

## Discussion

Current targeted therapies in cancer are designed against a handful of oncogenic proteins that drive disease progression. In an effort to expand the universe of potentially exploitable anti-cancer targets, recent genomic efforts have started to uncover therapeutic vulnerabilities arising from copy number losses not only affecting driver but passenger genes in cancer ^13,14^. However, it is not currently known whether genes collaterally affected by amplifications can drive similar sensitivities. In this study, we sought to define and characterize the class of ‘amplification-related gain of sensitivity’ (ARGOS) genes, causing a proliferation defect in cancer cells when overexpressed often due to their close proximity to focal oncogenic driver amplifications.

Gene expression upon amplification is generally considered dosage-sensitive ^4^, but previous studies have shown extensive compensation of protein expression ^5,40^. To measure gene expression compensation, studies have so far quantified the degree of overexpression with genomic amplifications ^12,40^. Here, we instead quantified the evidence for a deviation of gene expression from the expected dosage-sensitive scaling. We preferred using this approach, so that genes that scale with copy number but are noisy in their expression levels would not be falsely identified as compensated. Similarly, we preferred using mRNA over protein expression, as the available data are more complete and as RNA levels are more strongly correlated with DNA levels than proteins.

Our ARGOS genes represent a new class of sensitivities that would go unnoticed in more traditional efforts to identify dependency genes, such as RNA interference or CRISPR-Cas9 mediated knock-out screens. Hence, this joint computational and experimental approach enables the rational identification and prioritization of compensated genes with strong biological evidence for their toxicity. We provide the identified compensated, toxic, and ARGOS gene lists as a resource to the community, for follow-up in individual as well as across cancer types.

We selected *CDKN1A* and *RBM14* as proof-of-concept candidates to identify putative mechanisms of toxicity arising from genomic amplification. For this, we engineered cell line models where protein overexpression could be induced at levels equivalent to those achieved by copy number gains present in TCGA and CCLE samples. Whereas a strong toxicity and compensation profile could be expected for the tumor suppressor and cell cycle regulator CDKN1A, the effects of RBM14 overexpression on cancer cell growth remained unknown. Previous studies identified *RBM14* as an essential gene during mouse embryogenesis ^48^, with its genetic depletion leading to increased genomic instability and cell death by apoptosis. Additionally, RBM14 has been implicated in DNA repair by c-NHEJ ^46,60^, and its knock-out sensitizes glioblastoma cells to radiotherapy ^61^. Here, we find that overexpression of RBM14 in the range of gained copies can already cause a similarly detrimental effect: (1) gene overexpression leads to preferential DNA damage repair by c-NHEJ, an error-prone process that increases the rate of aberrant cell divisions; (2) upregulation of RBM14 and DNA-PK as members of the HDP-NRP complex induces cGAS-STING pathway activation.

In accordance with protein stoichiometry imbalances driving a common phenotype ^62^, the toxicity effects in our cell line models were observed regardless of RBM14 gene amplification status. Interestingly, the proliferation defect via pro-inflammatory signaling of RBM14 is in line with the strong enrichment for immune response pathways observed in genes that drop out in the analyzed ORF screens, supporting its putative role as a cell-intrinsic driver of toxicity. However, we do not expect RBM14’s mechanism of toxicity to be limited to DNA damage response and innate immune response regulation, as both our ORF screen toxicity analysis and initial validation of lung and breast cancer cell lines showed gene overexpression to strongly decrease cellular proliferation and protein translation rates in the absence of IR-mediated DNA damage. Identification of all biological processes associated with RBM14 overexpression, as well as their relative contribution, will require further investigation in additional cancer cell line models as well as in *in vivo* experiments.

Finally, our compensation and toxicity analyses identified 26 additional ARGOS genes that remain to be validated in pan-cancer (6) and lineage-specific (20) manners. However, this number may be an underestimation of the full universe of targets, as the ORF screens we analyzed were performed with lentiviral libraries that only span about half of the protein-coding genome (12,753 genes) and were conducted in a limited number of tumor types.

This study establishes a compendium of genes compensated across human cancers and in tissue-specific patterns, with functional evidence for toxicity effects associated with gene overexpression in the context of this disease. We predict the discovery of additional candidates as data from other functional genomic approaches (such as CRISPR activation screens) becomes more widely available. As ARGOS genes can constitute cellular liabilities, their discovery could ultimately lead to the emergence of therapeutic opportunities in tumors harboring ARGOS gene amplification.

## Acknowledgements

We thank the following investigators for providing data related to ORF screens: Cory Johannessen and Tikvah K. Hayes in Meljuso, OVSAHO and Kuramochi cell lines; Nikhil Wagle, Pingping Mao and Ofir Cohen for T47D cells; Adam Bass, Bruno Bockorny and Maria Rusan for NCI-H2077 cells. U2OS reporter cell lines were kindly provided by the Stark Laboratory and used in the analysis of DNA damage repair as described in ^56^. This work was funded by a Marie Skłodowska-Curie postdoctoral fellowship (101068734) to MSR and a Dutch Cancer Society grant (18-RUG-11457) to FF. Work in the U.B.-D. lab was supported by an ERC Starting Grant (StG #945674). KS was supported by NIH 1R35 CA283977-01. RB was supported by The Gray Matters Brain Cancer Foundation, Brain Tumour Charity, Break Through Cancer, and the Brown Fund for Innovative Cancer Informatics.

## Disclosures

KS is on the SAB and has stock options with Auron Therapeutics. KS receives grant funding from Novartis and KronosBio on topics unrelated to this manuscript. WCH is a consultant for Thermo Fischer Scientific, Solasta Ventures, MPM capital, KSQ Therapeutics, Tyra Biosciences, Frontier Medicines, Jubilant Therapeutics, RAPPTA Therapeutics, Serinus Biosciences, Hexagon Biosciences, Kestral Therapeutics, Function Oncology, and Calyx. RB consults for Scorpion Therapeutics and receives grant funding from Novartis.

## Methods

### Code availability

Unless stated otherwise, computational analyses have been performed using R 4.3.1 and the Snakemake workflow manager. All analysis code, as well as a list of used packages and their versions, is available at https://github.com/mschubert/ToxicGenes.

### Data sources

CCLE data was downloaded from DepMap (depmap.org) ^63^, including cell line annotations (“Cell_lines_annotations_20181226.txt”), RNA-seq gene counts (“CCLE_RNAseq_genes_counts_20180929.gct.gz”), and log_2_ copy number changes over the mean per gene (“CCLE_copynumber_byGene_2013-12-03.txt.gz”). Entrez gene IDs were remapped to HGNC symbols using Ensembl 104 ^64^. TCGA copy number call significance was obtained by GISTIC copy number calls, downloaded from the Broad TCGA copy number portal (tumorscape.org) using the Tumorscape 1.2.1 analysis with “2015-06-01 stddata 2015_04_02 arm-level peel-off” data ^1,65^. Genes were considered frequently amplified or deleted if they were identified by GISTIC in 15% or more of TCGA samples. Tumor purity estimates were used from the ESTIMATE algorithm ^66^ applied to the extended TCGA cohorts ^67^. Oncogenes and Tumor Suppressor gene lists were downloaded from the COSMIC gene census (version 96) ^36^ and included in the respective lists if they were listed as Hallmark, Tier 1, or Tier 2. Individual copy number events were extracted from previously published Ziggurat Deconstruction analysis of copy number states (“TCGA.all_cancers.150601.zigg_events.160923.txt”) ^65^.

### Gene compensation

The gene compensation analysis was performed either across tissues (pan-cancer) or for individual cancer types (tissue-specific), both for TCGA and CCLE data. Both CCLE and TCGA RNA-seq library size factors were calculated from raw counts using *DESeq2* (1.31.3) ^68^. Copy number counts were transformed from log_2_ into linear space and parameterized in euploid equivalents *c* and euploid deviation *d*. Therein, *c* represents the slope, *i.e.* the part of gene expression that is scaling with DNA copy number (0 if there are no DNA copies present, 1 if it is equal to the sample mean). *d* represents the deviation from this slope, with a value of 0 for the sample mean, − 1 if there are no DNA copies present, or 1 if there are twice the base copies. Hence, *c* is always one integer unit higher than *d*. The expression scaling term *c* is fitted per tissue *t*, allowing for different basal expression levels per tissue. This leaves us, for the CCLE, with the regression formula for each gene across samples:

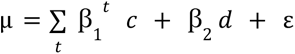

For the TCGA, we multiplied the cancer component of the gene expression by its purity term *p*, and add an additional term to represent the gene expression contribution of the non-cancer compartment:

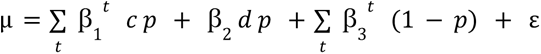

Both formulas are equivalent for a purity value of 1, as in this case ꞵ_3_ becomes zero and in the first two *p* terms can be dropped. Note that for TCGA *c* and *d* are DNA copies for cancer cells, hence *c p* and *d p* represent the observed DNA copy number in the mixture. Each µ represents the mean gene counts

observed for a given gene in RNA-seq data divided by library size. For each gene, we then fit a Negative Binomial regression:

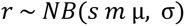

Here, for the total number of observed reads *r* and in addition to a theoretical gene mean µ we need to take into account the RNA-seq library size *s*, as well as the mean expression of a gene *m* (so we obtain comparable parameters independent of gene expression level and library size). We fit this regression using the R package *brms* (version 2.14) ^69^ using a tissue-specific scaling term (ꞵ_1_), a common compensation term ( ꞵ_2_), in the case of TCGA data a tissue-specific purity term (ꞵ_3_), and a linear link function for β. σ represents the shape parameter. We used the default prior for σ and set the following priors on *s m* µ ꞵ*_i_*:

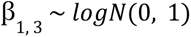

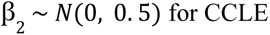

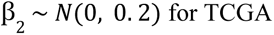

We included from the CCLE/TCGA all samples where a gene has no copy number changes (15% tolerance) and at least 3 or 5 samples with an amplification, respectively (1 copy number with 15% tolerance). We compared our proposed model with a model lacking a purity term (**Fig. S2a**, left) and a model with a common purity term across tissues (**Fig. S2a**, center), and in both cases found an inflation of compensation in the TCGA over the CCLE, which is why we did not use these models. As proposed, the result in TCGA and CCLE showed the strongest correlation without an obvious inflation effect in the TCGA (**Fig. S2a**, right). We normalized ꞵ_2_ by the mean euploid expression 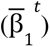 and calculated statistical significance using the z-score between the posterior and the origin of ꞵ_2_. To arrive at the final compensation score *s*, we shrank ꞵ_2_* by its pseudo-p-value derived from the z-score:

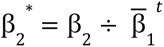

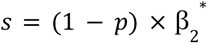

We estimated gene-set level differences using a linear model on the compensation scores of each gene in a set.

### Protein compensation

We used normalized protein expression data from CCLE proteomics ^70^ and reverse log_2_-transformed them back into a linear scale. We then estimated the expected scaling between no expression without DNA copies and observed average expression (1 for normalized data), as well as observed scaling using a linear regression model.

### ORF screen analysis

The ORF screen (“toxicity”) analysis was performed in both a pan-cancer and tissue-specific manner. We repurposed the vehicle treatment arm of 17 independent ORFeome library screens that were conducted to assess drug sensitivity across eight tumor types (Genetic Perturbation Platform, Broad Institute). For each screen, cells were infected with the ORFeome pLX317 barcoded library (16,100 barcoded ORFs overexpressing 12,753 genes), selected with puromycin and cultured for 14-21 days. We compared the high-throughput sequencing data from early (after puromycin selection) and late (after 14-21 days of cell culture) time points to estimate log_2_-fold changes and significance in ORF barcode representation using a linear model, where each barcode measurement for a given gene in any screen was considered an independent observation. All screens were included for the pan-cancer analysis, whereas the tissue-specific analysis was limited to screens that share a cancer type. We estimated gene-set level differences using a linear model on the Wald statistic of each gene in a set.

### Cell line information and tissue culture

NCI-H838 (cat. no. CRL-5844; ATCC), NCI-H1650 (cat. no. CRL-5883; ATCC), SK-LU-1 (cat. no. HTB-57; ATCC), ZR-75-1 (cat. no. CRL-1500; ATCC), HCC70 (cat. no. CRL-2315; ATCC), and MDA-MB-231 (cat. no. HTB-26; ATCC) cells were cultured in RPMI 1640 medium (Gibco) supplemented with 10% tetracycline negative fetal bovine serum (FBS; Gemini Bio) and 1% penicillin-streptomycin (Invitrogen). HEK-293T cells (cat. no. CRL-11268; ATCC) were grown in DMEM (Gibco) with 10% FBS. U2OS EJ7-GFP and DR-GFP cells were a gift from Dr. Jeremy Stark (City of Hope) and were cultured in McCoy’s 5A medium (Gibco), supplemented with 10% FBS and 1% penicillin/streptomycin. All cell lines were maintained at 37°C in 5% CO_2_ and frequently examined for mycoplasma contamination using the MycoAlert Mycoplasma Detection Kit (Lonza).

### Plasmid construction

Open reading frame (ORF) constructs were obtained from the Genetic Perturbation Platform (Broad Institute). Entry pDONR223 vectors for CDKN1A (clone ID ccsbBroadEn_00282), RBM14 (clone ID ccsbBroadEn_02429) and luciferase control (clone ID BRDN0000464768) were cloned into the pLXI-403 doxycycline-inducible destination vector (clone ID BRDN0000464768) by Gateway®-compatible LR clonase II (Invitrogen). This final expression vector contained the coding sequence of each gene followed by an in-frame V5-tag sequence. Following Reaction products were transformed in One Shot Stabl3 chemically competent *E. coli* (Invitrogen) under 100 μg mL^-1^ carbenicillin selection (Sigma-Aldrich). Plasmid DNA from single colonies was extracted with the QIAprep Spin Miniprep Kit and Maxi Kits (Qiagen) and confirmed by Sanger sequencing (Genewiz) using the primer 5’-GAC GTG AAG AAT GTG CGA GA-3’.

### Lentiviral transduction

Lentivirus-compatible expression vectors generated from Gateway Cloning were transfected in HEK-293T cells with Lipofectamine 3000 Transfection Reagent (cat. no. L3000001, Invitrogen) along with psPAX2 and pVSV-G viral packaging plasmids (Addgene) following manufacturer’s protocol. Virus was harvested 48h later with a 0.45-micron syringe filter. For transduction of viral plasmids, cells were seeded at a density of 2 million cells/well in a 12-well plate. Virus (400 µL) was added together with 5 ug/mL of polybrene (Sigma-Aldrich) and cells were centrifuged at 2,000 rpm for 2 h at 30°C. The next day, cells were selected with 1 µg/mL puromycin (Gibco).

### Real-Time quantitative PCR (RT-qPCR)

One million cells were treated with 0-500 ng/mL doxycycline for 5 h. Following treatment, RNA extraction was conducted using the RNeasy Mini Kit (cat. no. 74004, Qiagen) with an on-column DNase digestion (cat. no. 79254, Qiagen). Next, 2 µg of purified RNA was converted to cDNA with the Maxima H Minus first-strand cDNA synthesis kit (Thermo Fisher), using the random hexamer primer provided per manufacturer’s protocol. For RT-qPCR analysis, 500 ng of resulting cDNA was added to a reaction mixture containing 1X Maxima SYBR Green/ROX qPCR Master Mix (cat. no. K0221, Thermo Fisher Scientific) and 0.3 µM forward and reverse primers (IDT) against CDKN1A (forward 5’-GGA AGA CCA TGT GGA CCT GT-3’; reverse 5’-GGA TTA GGG CTT CCT CTT GG-3’), RBM14 (forward 5’-TTT TCG TGG GCA ATG TGT CGG C-3’; reverse 5’-GAT TGC GGC TTT GGC ATC TGC T-3’), and actin (forward 5’-CCG AAA GTT GCC TTT TAT GG-3’; reverse 5’-TCA TCA TCC ATG GTG AGC TG-3’). Samples were run in a StepOne RT-qPCR thermocycler, using the following running protocol: 95°C for 15 sec and 60°C for 1 min for 40 cycles, with melt curve stage at 95°C for 15 sec and 60°C for 1 min. Statistical analysis was performed by the ΔΔC_T_ method using the StepOne Software v2.1 (Applied Biosystems).

### Immunoblotting

Cells were treated with 0-500 ng/mL doxycycline (Takara Bio) 48 h prior sample collection. Pellets from 2 million cells were collected and lysed with RIPA lysis buffer (20 mM Tris-HCl at pH 7.5, 150 nM NaCl, 1 mM Na_2_EDTA, 1 mM EGTA, 1% NP-40, 1% sodium deoxycholate, 2.5 mM sodium pyrophosphate, 1 mM beta-glycerophosphate, 1 mM Na_3_VO_4_, 1 μg/mL leupeptin), phenylmethanesulfonyl fluoride (PMSF; Cell Signaling Technology) and a protease and phosphatase inhibitor cocktail (Boston BioProducts). Lysates were incubated for 45 min on ice with pulse-vortexing in 15 min intervals and centrifuged at 13,000 rpm for 10 min in 4°C. Protein concentrations were quantified by BCA against a bovine serum albumin standard (ThermoFisher). Absorbance was measured by incubating protein samples and BCA standards with Pierce A/B colorimetric reagents (Thermo Scientific). A total of 25 µg for each sample was loaded on a NuPAGE 4-12% Bis-Tris gel with 1X NuPAGE MOPS Running buffer (Life Technologies, Invitrogen) and separated for 1.5 h at 120V. For protein size comparison, the Cytiva full-range Rainbow^TM^ molecular weight marker (Fisher Scientific) was used as reference. Following protein separation, a dry transfer was completed at 30V for 6 min with PVDF stacks in an iBlot 2 transfer instrument (Thermo Fisher). After gel transfer, the membrane was blocked in 5% dry milk in TBS-T for 30 min and incubated overnight at 4°C with primary antibodies against: rabbit V5 at 1:1,000 (Cell Signaling Technology), mouse vinculin at 1:1,000 (Millipore Sigma), rabbit CDKN1A at 1:1,000 (Cell Signaling Technology), and rabbit RBM14 at 1:1,000 (Abcam) in blocking solution containing 5% milk and TBS-T. For the detection of phospho-DNA-PKcs, samples were run in a 3-8% Tris-acetate gel with 1X NuPAGE MES Running buffer (Life Technologies, Invitrogen). A wet transfer was done overnight at 30 V, and the membrane was blocked in 5% BSA in TBS-T. Overnight incubation was done with primary antibodies against rabbit DNA-PKcs S2056 (Cell Signaling Technology) and mouse vinculin at 1:1,000 (Millipore Sigma). After incubation with primary antibodies, the membranes were washed in TBS-T and further incubated for 1 h with goat anti-rabbit at 1:3,000 (ThermoFisher) and goat anti-mouse (ThermoFisher) at 1:10,000 dilution. Signal was detected by chemiluminescence with SuperSignal West Pico and Femto kits (Thermo Scientific) in ImageQuant LAS 4000 imager (GE Healthcare Life Sciences). Blots were quantified using ImageJ 1.52 k.

### Luciferase reporter assay

The expression of inducible luciferase in cell lines was measured using the Dual-Glo Luciferase Assay system (cat. no. E2920, Promega). Cells were seeded at a density of 15,000 cells per well in a 96-well plate (3917, Costar). The following day, 500 ng/mL doxycycline (Takara Bio) was added to the media and cells were incubated for 48 h at 37°C. Luciferase buffer and substrate reagents were prepared per manufacturer’s instruction. Briefly, the Dual-Glo Luciferase reagent was added to a 1:1 ratio to the cells and allowed to incubate at room temperature (25°C) for 10 minutes in the darkness on an orbital shaker. Firefly luciferase expression (luminescence) was measured using the SpectraMax M5 plate reader and Dual-Glo luciferase protocol with an integration time of 500 ms.

### Growth curves

Cells were seeded in 6 technical replicates at a density of 5,000 cells per well in a 96-well plate. The next day, cells were treated with 0-500 ng/mL doxycycline (Takara Bio) and imaged in an Incucyte Live-Cell Analysis system (Sartorius) to track confluency over the course of 7 days. Images were acquired with a 10X objective every 6 hours, and analyzed with custom-defined masks to identify the area of each cell line. Media was replaced with fresh doxycycline every 48 h.

### Nascent Protein Synthesis Assay

Nascent protein synthesis was measured with the Click-iT HPG Alexa Fluor 488 Protein Synthesis Assay Kit (cat. no. C10428, ThermoFisher). Cells were plated in 12-well plates (Cell Treat) containing 18 mm cover glasses (Fisher Scientific) at the bottom, and treated with 500 ng/mL of doxycycline (Takara Bio) for 48 h. Following induction, cells were washed with PBS and incubated with 50 M of Click-iT homopropargylglycine (HPG) in methionine-free RPMI media (Gibco) for 30 min. Next, cells were fixed with 3.7% formaldehyde in PBS and permeabilized using 0.5% Triton X-100. The Click-iT reaction was performed per manufacturer’s protocol by preparing a reaction cocktail mix containing Alexa Fluor 488 azide. DNA staining was performed using NuclearMask Blue Stain, and incubated for 30 min in the dark. Coverslips were then removed from the plate and mounted on glass slides using Slow-Fade mounting solution (Fisher Scientific). Slides were imaged in an Olympus IX73 inverted microscope using the CellSens Software (Olympus). GFP signal intensity was quantified and normalized to the total number of cells using Image J.

### Cell cycle analysis by flow-cytometry

Cell cycle analysis was performed by *in vitro* labeling of cells with the APC BrdU Flow Kit (cat. no. 552598, BD Biosciences). First, 1 million cells were treated with 500 ng/mL doxycycline for 48 h. Next, 10 μM BrdU solution was added to the culture media and cells were incubated for 2 h. Cells were then fixed and permeabilized according to the manufacturer’s protocol. Next, samples were treated with 60 μg of DNase and incubated for 1 h at 37 ^॰^C to expose the incorporated BrdU. Finally, cells were stained for BrdU (1:50 antibody dilution) and total DNA (7-AAD 1:50 solution dilution) and analyzed by flow cytometer in a BD LSRFortessa cell analyzer (BD Biosciences) using 488 nm and 640 nm lasers. Cells were gated and assigned to G1, S, or G2 phases of the cell cycle.

### Quantification of apoptosis by flow-cytometry

Apoptotic cells were labeled for flow cytometry using a Dead Cell Apoptosis Kit with Annexin V FITC & Propidium Iodide (cat. no. V13242, Invitrogen). Briefly, cells were seeded at a density of 500,000 cells/well in a 6-well plate and treated with 500 ng/mL doxycycline for 72 h. Next, cell pellets were collected and resuspended in 100 µL of 1X annexin-binding buffer containing 5 µL of FITC annexin V antibody and 1 µL of 100 µg/mL propidium iodide (PI) working solution. Following an incubation at room temperature for 15 min, 400 µL of 1X annexin-binding buffer was added to each sample. Stained cells were analyzed by flow cytometry in a BD LSRFortessa cell analyzer (BD Biosciences), measuring the fluorescence emission at 530 nm (FITC) and 575 nm (PI). Cells were gated and classified as live (negative for both annexin V and PI), early apoptotic (positive for annexin V), or late apoptotic (positive for both annexin V and PI). Image quantification was performed in FlowJo v 10.8.1 software.

### Immunofluorescence analysis

Cells were seeded at a density of 40,000 cells per well in a 12-well plate with round coverslips. After 48h treatment with 500 ng/mL doxycycline, cells were subjected to 2 Gy X-ray ionizing radiation (IR). At each time point (0, 15, 30, 60, 120 and 360 min post-radiation), cells were washed with CSK buffer for 5 min and later incubated with 0.7% Triton X-100 in CSK for 5 min on ice. Fixation was done with 4% paraformaldehyde in CSK at room temperature for 30 min. After two washes with PBS, cells were permeabilized with 0.2% Triton X-100 in PBS and blocked with PBS-T containing 5% BSA for 1 h. Incubation with 1:3000 or 1:100 dilution primary antibodies against mouse γH2A.X (cat. no. 05-636, Millipore) or STING (cat. no. PA5-23381, Invitrogen) was done overnight at 4°C. After 3 washes with PBS, cells were incubated with fluorescent AF-568 goat anti-mouse secondary antibody (cat. no A11031, Invitrogen) and Hoechst 33342 nucleic acid stain (cat. no. H3570, Thermo Fisher) at a 1:10,000 dilution for 1 h in the dark. Coverslips were mounted on glass slides by addition of SlowFade Diamond antifade mountant solution (cat. no. S36963, ThermoFisher), and visualized either in an Olympus IX73 Inverted Microscope (γH2A.X immunofluorescence) or in a SP8 TCS Leica confocal microscope (STING immunofluorescence). Image processing, quantification of γH2AX foci per nucleus, and quantification of STING area was performed in ImageJ2 v 2.9.0.

### Double-strand-break repair GFP reporters

DNA repair pathway choice was measured in U2OS EJ7-GFP (c-NHEJ) and U2OS DR-GFP (HR) cells as described in ^56^. Cells were seeded at a density of 500,000 cells/well in 12-well plates. The next day, cells were transfected with 500 ng pCAGGS-NZEGFP (transfection efficiency control), 500 ng pCAGGS-BSKX (empty vector control), 500 ng pCBASce (ISceI; for HR assay) and 250 ng pX330-7a/ 250 ng pX330-7b (‘7a’ and ‘7b’ guide RNAs; for c-NHEJ assay) plasmids using the Lipofectamine 3000 Transfection Reagent (cat. no. L3000001, Invitrogen). After 8 h, fresh media was added with 500 ng/mL doxycycline. After 48 h, cells were collected, resuspended in 500 µL eBioscience flow cytometry staining buffer (cat. no. 00-4222-26, Invitrogen) and analyzed in a BD LSRFortessa cell analyzer (BD Biosciences), measuring the fluorescence emission at 530 nm (GFP). GFP-positive cells were quantified in each condition with FlowJo v 10.8.1 software.

### Drug dose-response curves

Cells were seeded at a density of 10,000 cells per well in a 96-well plate and treated with 500 ng/mL doxycycline. The next day, cells were treated with 0-100 µM of the DNA-PK inhibitor NU7441 (cat. no. HY-11006, MedChemExpress) or RAD51 inhibitor B02 (cat. no. HY-101462, MedChemExpress) and incubated for 72 h. The CellTiter-Glo 2.0 viability assay (cat. no. G9241, Promega) was used to measure the level of ATP as a surrogate for cell viability. Luminescence was measured in a SpectraMax M5 plate reader (Associated Technologies Group) with the SoftMax Pro software using the CellTiter-Glo protocol with an integration time of 500 ms. 0

### Time-lapse imaging of mitosis

PIGPZ–H2B–Cherry was subcloned from an existing H2B–Cherry construct using *NotI* and *BsrGI* sites. Lentiviral transduction of pIGZ H2B–Cherry was achieved by transfecting 3 μg of the desired plasmid supplemented with the following packaging plasmids: 3 μg pSPAX2 and 1 μg pMD2.G into HEK-293T cells. Plasmids psPAX2 (Addgene plasmid, 12260) and pMD2.G (Addgene plasmid, 12259) were gifts from Prof. D. Trono (École Polytechnique Fédérale de Lausanne). After 48 h, medium was collected, passed through a 0.45-μm filter (VWR Science), and transferred onto the target cells. To increase transduction efficiency, polybrene (Millipore) was added to a final concentration of 8 μg/ mL. For time-lapse imaging, 60,000 cells expressing H2B–mCherry were first seeded per well on imaging chambers (LabTek) and treated with 2 Gy IR and immediately replaced with fresh medium. All cells were subsequently imaged for 8 hours on a DeltaVision Elite imaging station (GE Healthcare), equipped with a CoolSNAP HQ2 camera, a 40X 0.6 NA immersion objective (Olympus), and DeltaVision softWoRx software. Images were acquired at 6 min intervals and included z-stacks of 20 images at 0.4 μm intervals. Image analysis was done using ICY software (Institut Pasteur). Only cells that entered mitosis and stayed in frame throughout the imaging session were included for analysis.

## References

1. Beroukhim, R. et al. The landscape of somatic copy-number alteration across human cancers. Nature 463, 899–905 (2010).

2. Zack, T. I. et al. Pan-cancer patterns of somatic copy number alteration. Nat. Genet. 45, 1134–1140 (2013).

3. Ben-David, U. & Amon, A. Context is everything: aneuploidy in cancer. Nat. Rev. Genet. 21, 44–62 (2020).

4. Fehrmann, R. S. N. et al. Gene expression analysis identifies global gene dosage sensitivity in cancer. Nat. Genet. 47, 115–125 (2015).

5. Gonçalves, E. et al. Widespread Post-transcriptional Attenuation of Genomic Copy-Number Variation in Cancer. Cell Syst 5, 386–398.e4 (2017).

6. Inaki, K. et al. Systems consequences of amplicon formation in human breast cancer. Genome Res. 24, 1559–1571 (2014).

7. Maruvka, Y. E. et al. Analysis of somatic microsatellite indels identifies driver events in human tumors. Nat. Biotechnol. (2017) doi:10.1038/nbt.3966.

8. Bhattacharya, A. et al. Transcriptional effects of copy number alterations in a large set of human cancers. Nat. Commun. 11, 715 (2020).

9. Schukken, K. M. & Sheltzer, J. M. Extensive protein dosage compensation in aneuploid human cancers. bioRxiv 2021.06.18.449005 (2021) doi:10.1101/2021.06.18.449005.

10. Ippolito, M. R. et al. Increased RNA and protein degradation is required for counteracting transcriptional burden and proteotoxic stress in human aneuploid cells. bioRxiv 2023.01.27.525826 (2023) doi:10.1101/2023.01.27.525826.

11. Alfieri, F., Caravagna, G. & Schaefer, M. H. Cancer genomes tolerate deleterious coding mutations through somatic copy number amplifications of wild-type regions. Nat. Commun. 14, 3594 (2023).

12. Mohanty, V., Wang, F., Mills, G. B., CTD2 Research Network & Chen, K. Uncoupling of gene expression from copy number presents therapeutic opportunities in aneuploid cancers. Cell Rep Med 2, 100349 (2021).

13. Nijhawan, D. et al. Cancer vulnerabilities unveiled by genomic loss. Cell 150, 842–854 (2012).

14. Muller, F. L. et al. Passenger deletions generate therapeutic vulnerabilities in cancer. Nature 488, 337–342 (2012).

15. Dey, P. et al. Genomic deletion of malic enzyme 2 confers collateral lethality in pancreatic cancer. Nature 542, 119–123 (2017).

16. Kryukov, G. V. et al. MTAP deletion confers enhanced dependency on the PRMT5 arginine methyltransferase in cancer cells. Science 351, 1214–1218 (2016).

17. Mavrakis, K. J. et al. Disordered methionine metabolism in MTAP/CDKN2A-deleted cancers leads to dependence on PRMT5. Science 351, 1208–1213 (2016).

18. Rendo, V. et al. Exploiting loss of heterozygosity for allele-selective colorectal cancer chemotherapy. Nat. Commun. 11, 1308 (2020).

19. Nichols, C. A. et al. Loss of heterozygosity of essential genes represents a widespread class of potential cancer vulnerabilities. Nat. Commun. 11, 2517 (2020).

20. Serrano, M., Lin, A. W., McCurrach, M. E., Beach, D. & Lowe, S. W. Oncogenic ras provokes premature cell senescence associated with accumulation of p53 and p16INK4a. Cell 88, 593–602 (1997).

21. Yaswen, P. & Campisi, J. Oncogene-induced senescence pathways weave an intricate tapestry. Cell 128, 233–234 (2007).

22. Liu, X.-L., Ding, J. & Meng, L.-H. Oncogene-induced senescence: a double edged sword in cancer. Acta Pharmacol. Sin. 39, 1553–1558 (2018).

23. Foijer, F. et al. Chromosome instability induced by Mps1 and p53 mutation generates aggressive lymphomas exhibiting aneuploidy-induced stress. Proc. Natl. Acad. Sci. U. S. A. 111, 13427–13432 (2014).

24. Hintzen, D. C. et al. The impact of monosomies, trisomies and segmental aneuploidies on chromosomal stability. PLoS One 17, e0268579 (2022).

25. Johannessen, C. M. et al. A melanocyte lineage program confers resistance to MAP kinase pathway inhibition. Nature 504, 138–142 (2013).

26. Bockorny, B. et al. RAS-MAPK Reactivation Facilitates Acquired Resistance in FGFR1-Amplified Lung Cancer and Underlies a Rationale for Upfront FGFR-MEK Blockade. Mol. Cancer Ther. 17, 1526–1539 (2018).

27. Iniguez, A. B. et al. Resistance to Epigenetic-Targeted Therapy Engenders Tumor Cell Vulnerabilities Associated with Enhancer Remodeling. Cancer Cell 34, 922–938.e7 (2018).

28. Bandopadhayay, P. et al. Neuronal differentiation and cell-cycle programs mediate response to BET-bromodomain inhibition in MYC-driven medulloblastoma. Nat. Commun. 10, 2400 (2019).

29. Guenther, L. M. et al. A Combination CDK4/6 and IGF1R Inhibitor Strategy for Ewing Sarcoma. Clin. Cancer Res. 25, 1343–1357 (2019).

30. Hayes, T. K. et al. A Functional Landscape of Resistance to MEK1/2 and CDK4/6 Inhibition in NRAS-Mutant Melanoma. Cancer Res. 79, 2352–2366 (2019).

31. Hwang, J. H. et al. CREB5 Promotes Resistance to Androgen-Receptor Antagonists and Androgen Deprivation in Prostate Cancer. Cell Rep. 29, 2355–2370.e6 (2019).

32. Stover, E. H. et al. Pooled Genomic Screens Identify Anti-apoptotic Genes as Targetable Mediators of Chemotherapy Resistance in Ovarian Cancer. Mol. Cancer Res. 17, 2281–2293 (2019).

33. Mao, P. et al. Acquired FGFR and FGF Alterations Confer Resistance to Estrogen Receptor (ER) Targeted Therapy in ER+ Metastatic Breast Cancer. Clin. Cancer Res. 26, 5974–5989 (2020).

34. Ghandi, M. et al. Next-generation characterization of the Cancer Cell Line Encyclopedia. Nature (2019) doi:10.1038/s41586-019-1186-3.

35. Weinstein, J. N. et al. The Cancer Genome Atlas Pan-Cancer analysis project. Nat. Genet. 45, 1113–1120 (2013).

36. Forbes, S. A. et al. COSMIC: exploring the world’s knowledge of somatic mutations in human cancer. Nucleic Acids Res. 43, D805–11 (2015).

37. Zerbib, J. et al. Human aneuploid cells depend on the RAF/MEK/ERK pathway for overcoming increased DNA damage. bioRxiv 2023.01.27.525822 (2023) doi:10.1101/2023.01.27.525822.

38. Collins, R. L. et al. A cross-disorder dosage sensitivity map of the human genome. Cell 185, 3041–3055.e25 (2022).

39. Yang, X. et al. A public genome-scale lentiviral expression library of human ORFs. Nat. Methods 8, 659–661 (2011).

40. Schukken, K. M. & Sheltzer, J. M. Extensive protein dosage compensation in aneuploid human cancers. Genome Res. 32, 1254–1270 (2022).

41. Sondka, Z. et al. The COSMIC Cancer Gene Census: describing genetic dysfunction across all human cancers. Nat. Rev. Cancer 18, 696–705 (2018).

42. Xiong, Y. et al. p21 is a universal inhibitor of cyclin kinases. Nature 366, 701–704 (1993).

43. Fredericks, A. M., Cygan, K. J., Brown, B. A. & Fairbrother, W. G. RNA-Binding Proteins: Splicing Factors and Disease. Biomolecules 5, 893–909 (2015).

44. Moore, K. S. & von Lindern, M. RNA Binding Proteins and Regulation of mRNA Translation in Erythropoiesis. Front. Physiol. 9, 910 (2018).

45. Zhou, L.-T. et al. A novel role of fragile X mental retardation protein in pre-mRNA alternative splicing through RNA-binding protein 14. Neuroscience 349, 64–75 (2017).

46. Simon, N. E., Yuan, M. & Kai, M. RNA-binding protein RBM14 regulates dissociation and association of non-homologous end joining proteins. Cell Cycle 16, 1175–1180 (2017).

47. Morchikh, M. et al. HEXIM1 and NEAT1 Long Non-coding RNA Form a Multi-subunit Complex that Regulates DNA-Mediated Innate Immune Response. Mol. Cell 67, 387–399.e5 (2017).

48. Li, J. et al. Rbm14 maintains the integrity of genomic DNA during early mouse embryogenesis via mediating alternative splicing. Cell Prolif. 53, e12724 (2020).

49. Shiratsuchi, G., Takaoka, K., Ashikawa, T., Hamada, H. & Kitagawa, D. RBM14 prevents assembly of centriolar protein complexes and maintains mitotic spindle integrity. EMBO J. 34, 97–114 (2015).

50. Fox, A. H. et al. Paraspeckles: a novel nuclear domain. Curr. Biol. 12, 13–25 (2002).

51. Davis, A. J., Chen, B. P. C. & Chen, D. J. DNA-PK: a dynamic enzyme in a versatile DSB repair pathway. DNA Repair 17, 21–29 (2014).

52. Mao, Z., Bozzella, M., Seluanov, A. & Gorbunova, V. Comparison of nonhomologous end joining and homologous recombination in human cells. DNA Repair 7, 1765–1771 (2008).

53. Brandsma, I. & Gent, D. C. Pathway choice in DNA double strand break repair: observations of a balancing act. Genome Integr. 3, 9 (2012).

54. Chan, D. W. & Lees-Miller, S. P. The DNA-dependent Protein Kinase Is Inactivated by Autophosphorylation of the Catalytic Subunit (*). J. Biol. Chem. 271, 8936–8941 (1996).

55. Bhargava, R. et al. C-NHEJ without indels is robust and requires synergistic function of distinct XLF domains. Nat. Commun. 9, 2484 (2018).

56. Gunn, A. & Stark, J. M. I-SceI-Based Assays to Examine Distinct Repair Outcomes of Mammalian Chromosomal Double Strand Breaks. in DNA Repair Protocols (ed. Bjergbæk, L.) 379–391 (Humana Press, 2012). doi:10.1007/978-1-61779-998-3_27.

57. Li, T. & Chen, Z. J. The cGAS-cGAMP-STING pathway connects DNA damage to inflammation, senescence, and cancer. J. Exp. Med. 215, 1287–1299 (2018).

58. Decout, A., Katz, J. D., Venkatraman, S. & Ablasser, A. The cGAS-STING pathway as a therapeutic target in inflammatory diseases. Nat. Rev. Immunol. 21, 548–569 (2021).

59. Mukai, K. et al. Activation of STING requires palmitoylation at the Golgi. Nat. Commun. 7, 11932 (2016).

60. Jang, Y. et al. Intrinsically disordered protein RBM14 plays a role in generation of RNA:DNA hybrids at double-strand break sites. Proc. Natl. Acad. Sci. U. S. A. 117, 5329–5338 (2020).

61. Yuan, M., Eberhart, C. G. & Kai, M. RNA binding protein RBM14 promotes radio-resistance in glioblastoma by regulating DNA repair and cell differentiation. Oncotarget 5, 2820–2826 (2014).

62. Brennan, C. M. et al. Protein aggregation mediates stoichiometry of protein complexes in aneuploid cells. Genes Dev. 33, 1031–1047 (2019).

63. Tsherniak, A. et al. Defining a Cancer Dependency Map. Cell 170, 564–576.e16 (2017).

64. Yates, A. D. et al. Ensembl 2020. Nucleic Acids Res. 48, D682–D688 (2020).

65. Mermel, C. H. et al. GISTIC2.0 facilitates sensitive and confident localization of the targets of focal somatic copy-number alteration in human cancers. Genome Biol. 12, R41 (2011).

66. Yoshihara, K. et al. Inferring tumour purity and stromal and immune cell admixture from expression data. Nat. Commun. 4, 2612 (2013).

67. Aran, D., Sirota, M. & Butte, A. J. Systematic pan-cancer analysis of tumour purity. Nat. Commun. 6, 8971 (2015).

68. Love, M., Anders, S. & Huber, W. Differential analysis of count data–the DESeq2 package. Genome Biol. (2014).

69. Bürkner, P.-C. Brms: An R package for Bayesian multilevel models using Stan. J. Stat. Softw. 80, 1–28 (2017).

70. Nusinow, D. P. et al. Quantitative Proteomics of the Cancer Cell Line Encyclopedia. Cell 180, 387–402.e16 (2020).

