## Supplementary figures and images for "A compendium of Amplification-Related Gain Of Sensitivity (ARGOS) genes in human cancer"

### Supplementary Figure 1

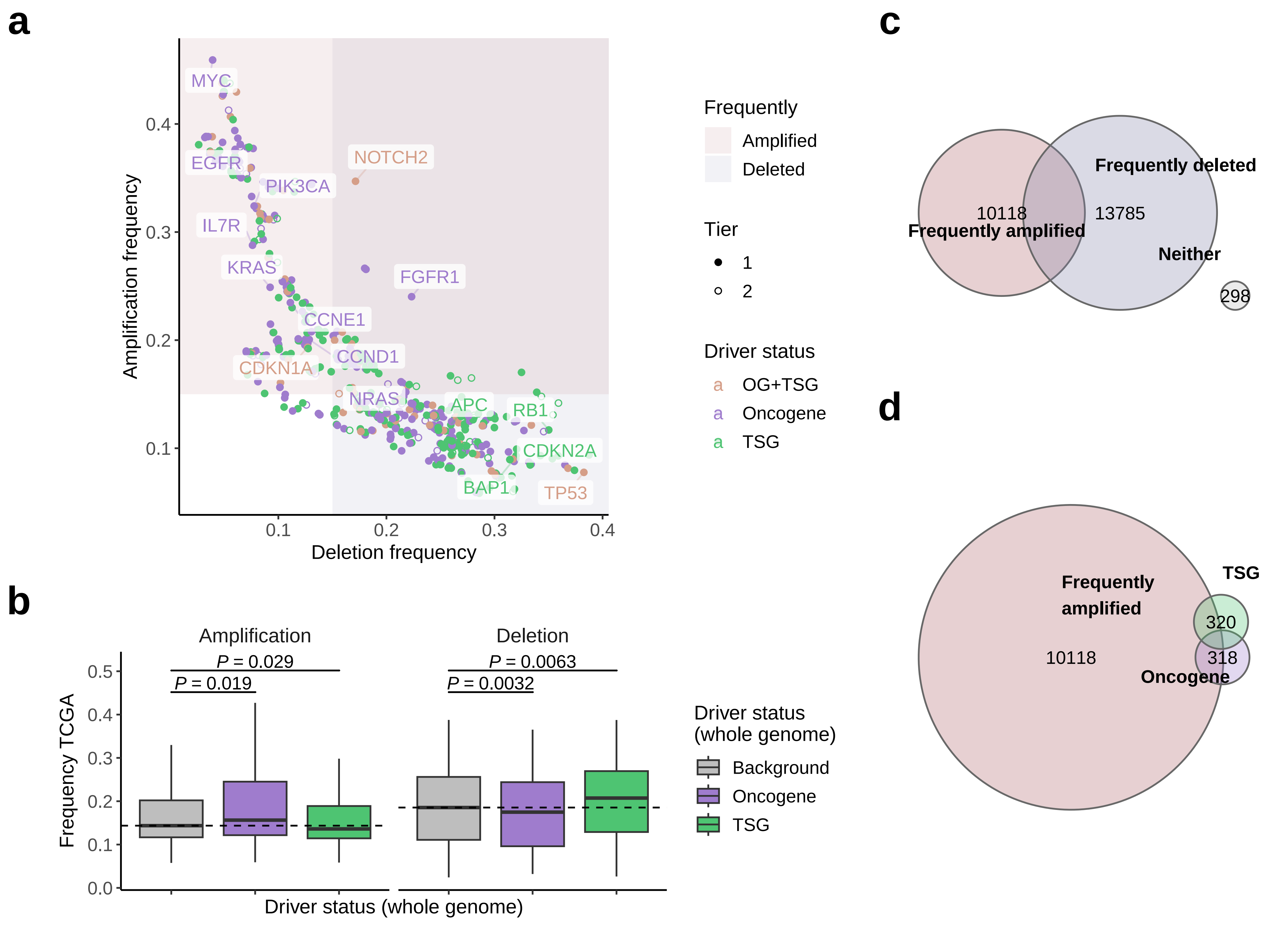

### Supplementary Figure 2

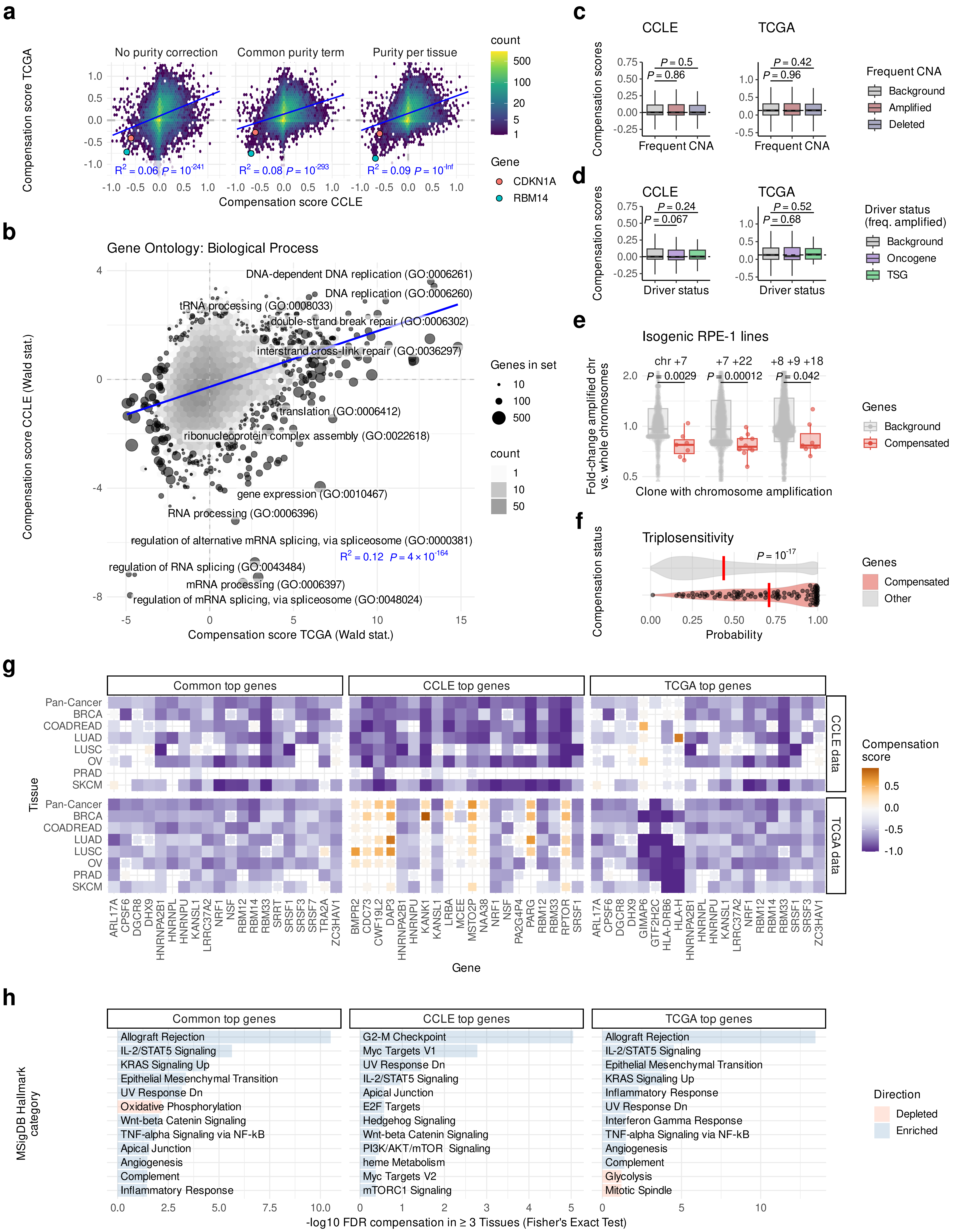

### Supplementary Figure 3

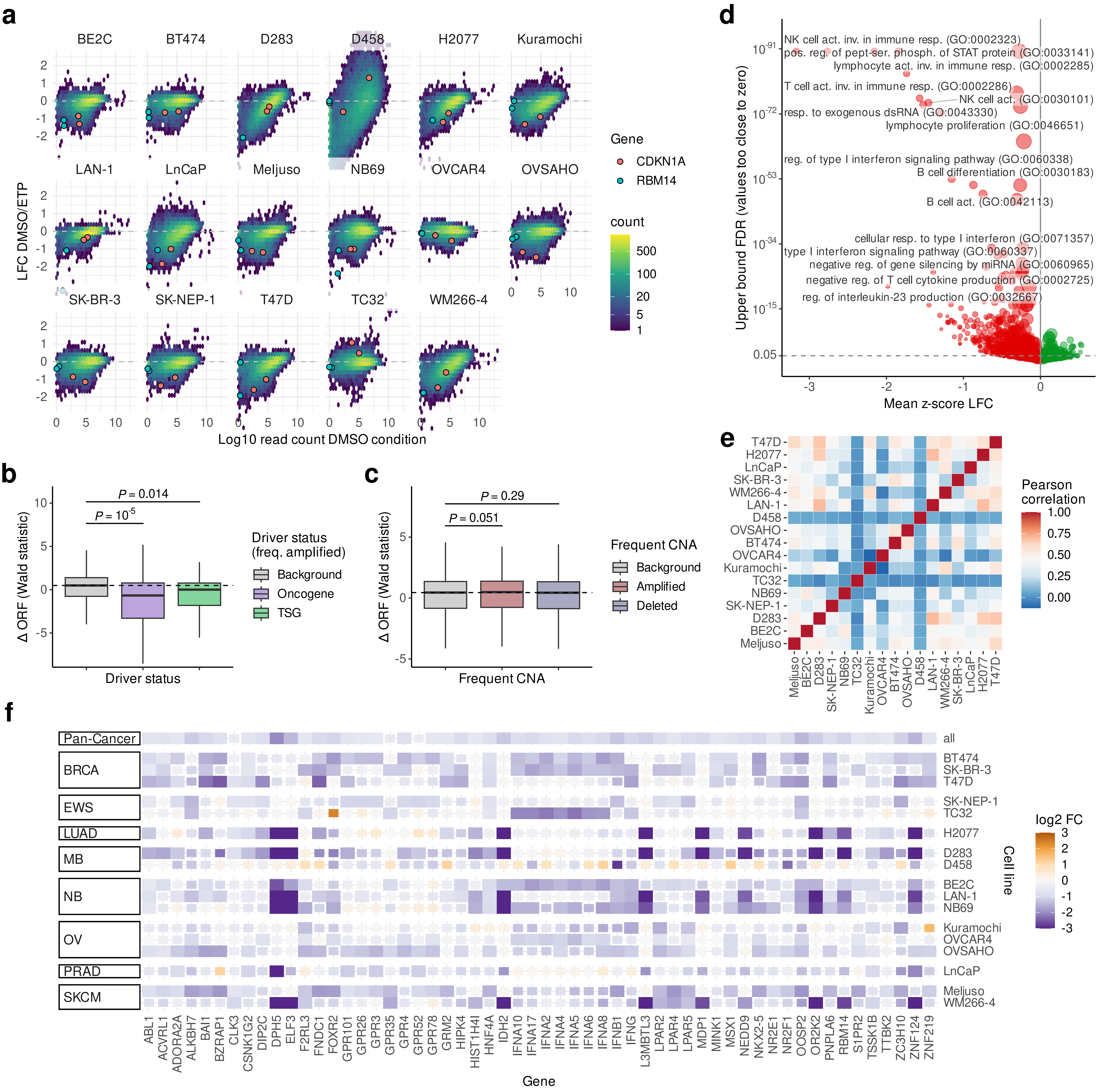

### Supplementary Figure 4

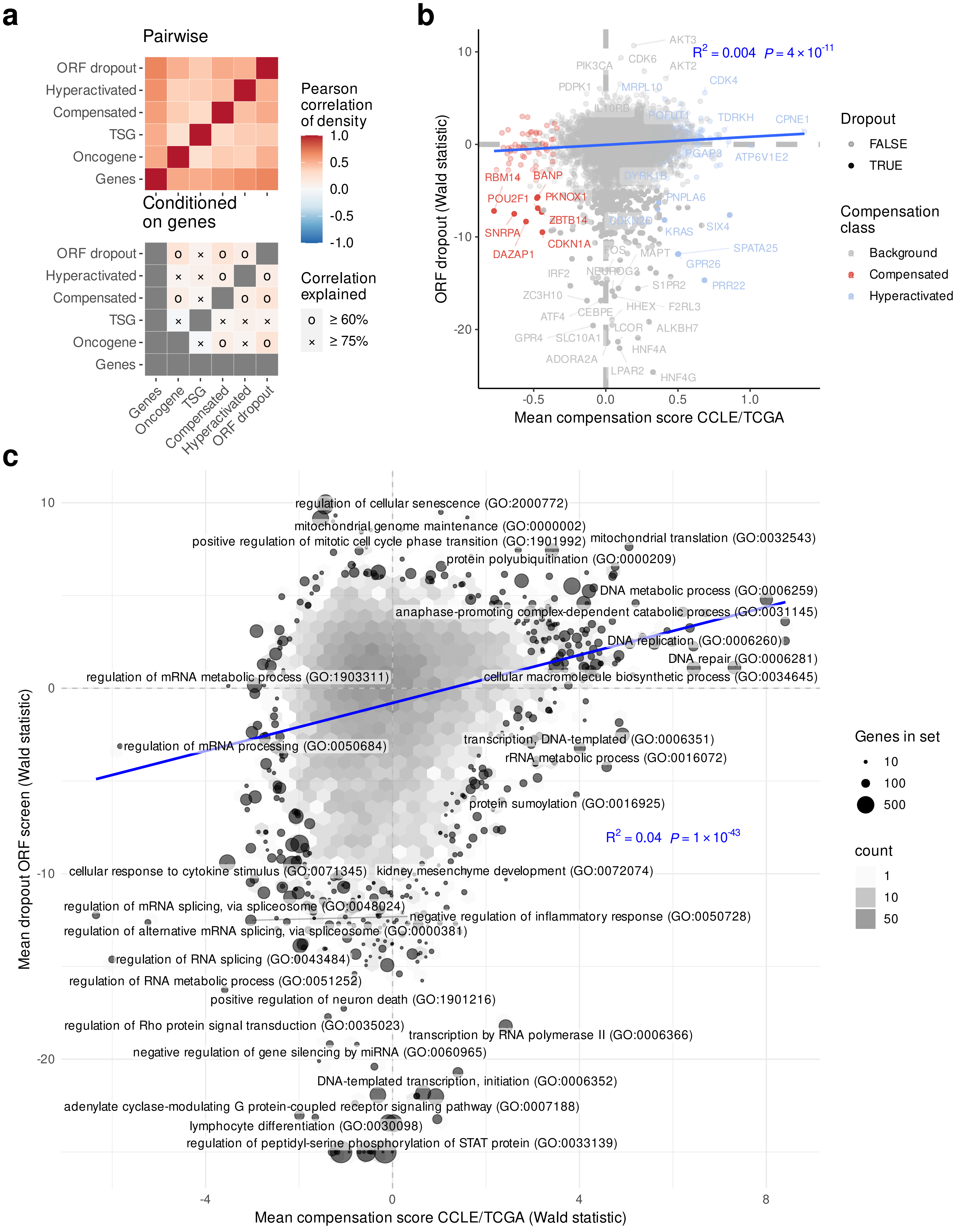

### Supplementary Figure 5

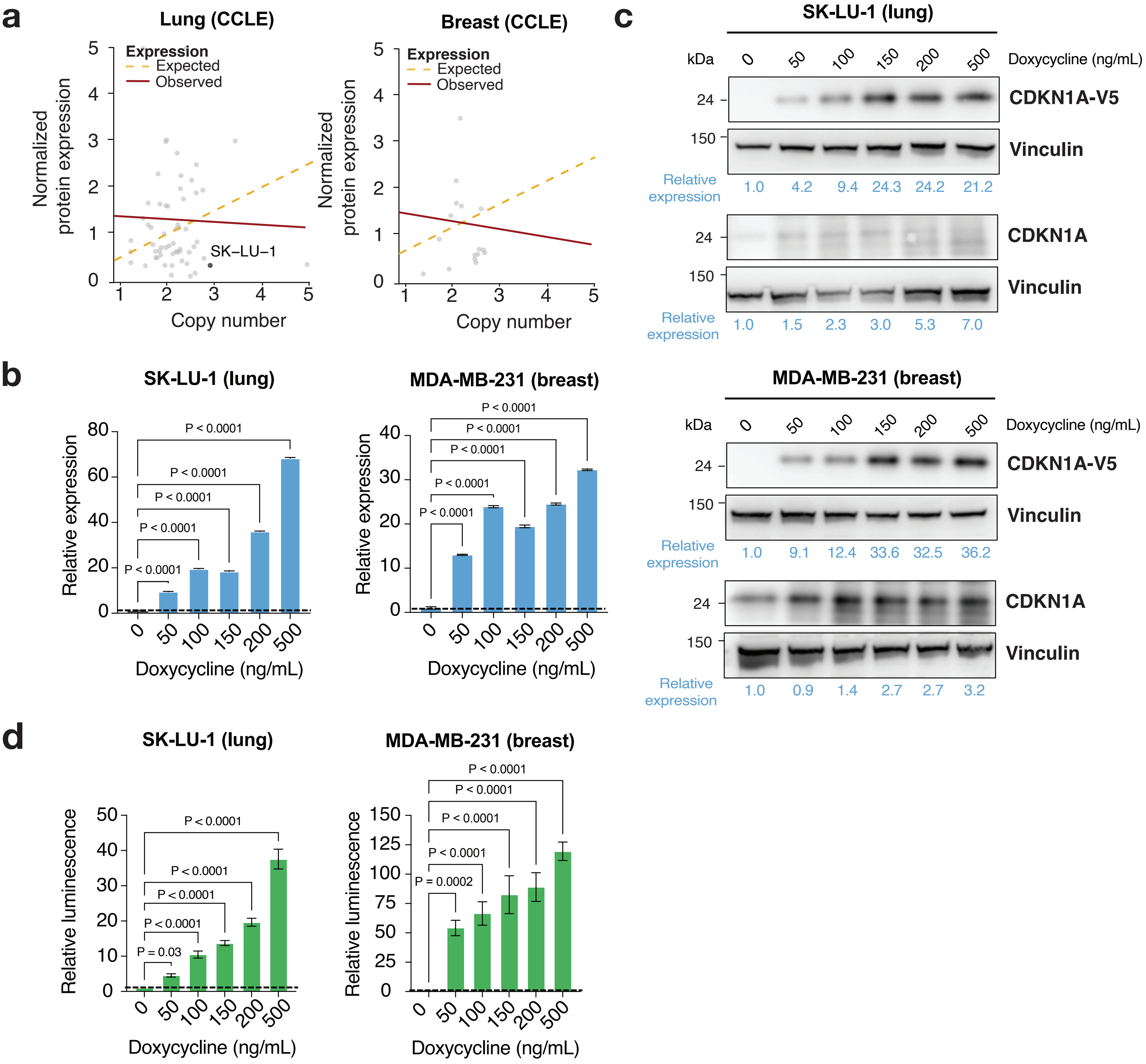

### Supplementary Figure 6

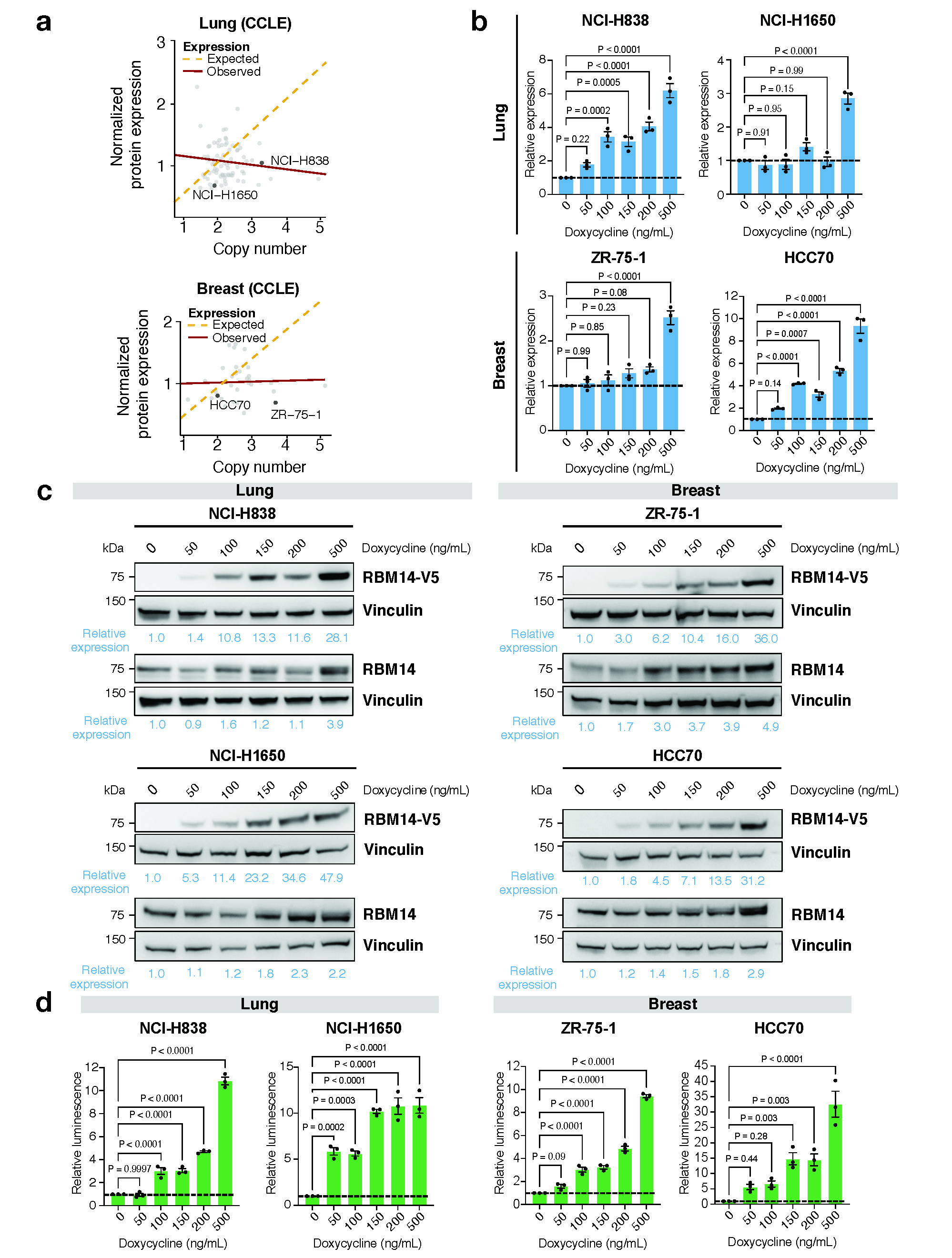

### Supplementary Figure 7

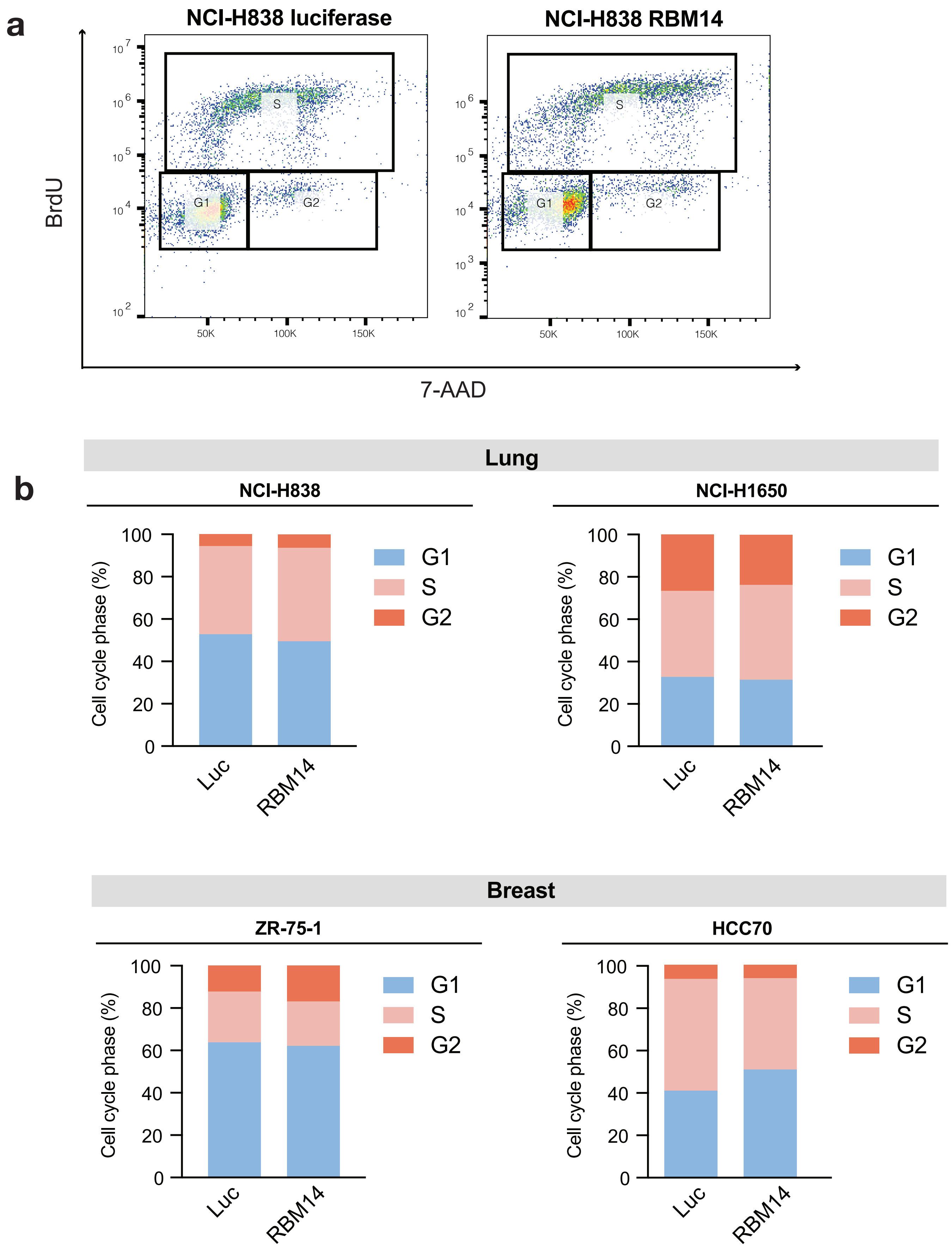

### Supplementary Figure 8

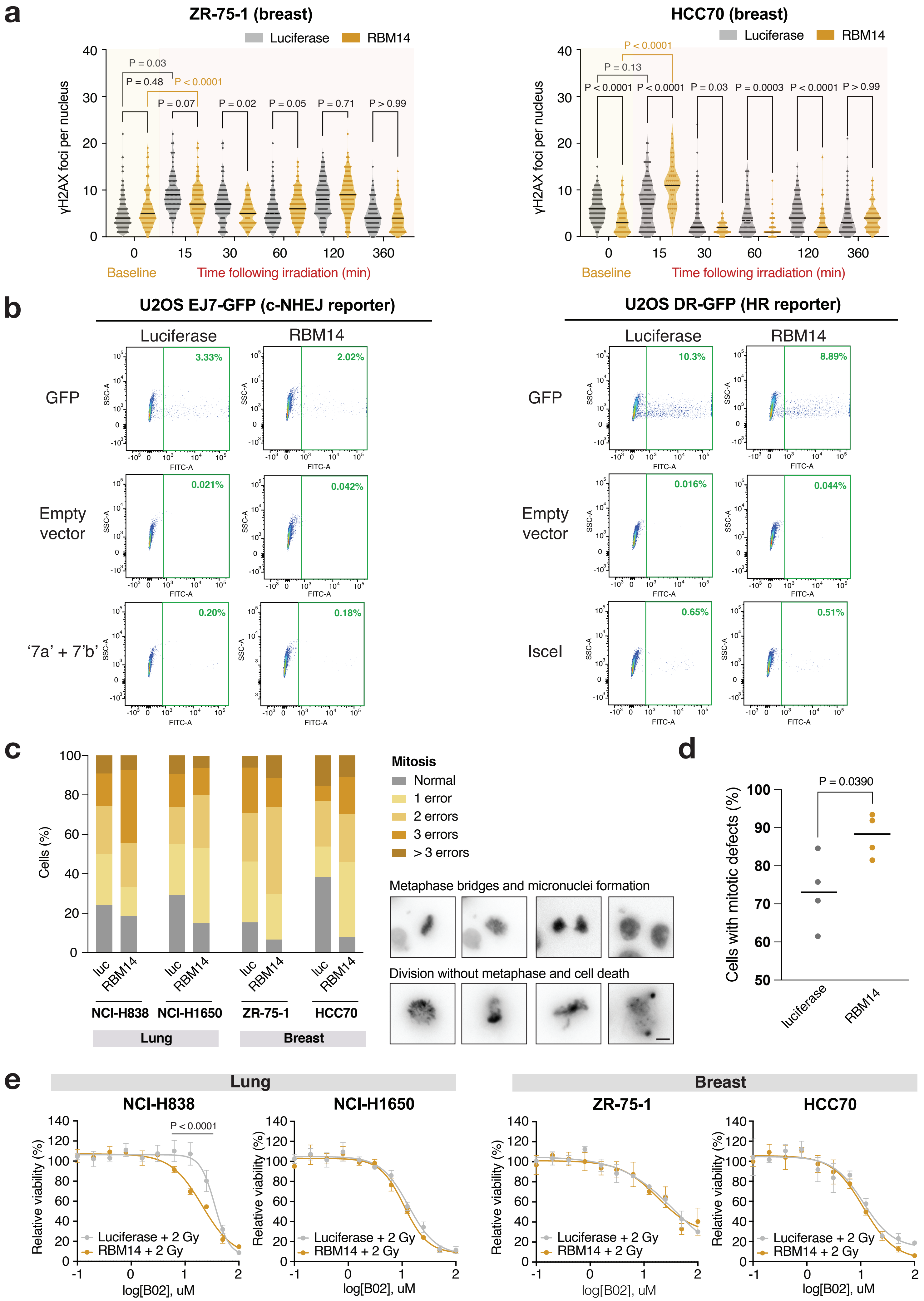
